# Probing different paradigms of morphine withdrawal on sleep behavior in male and female C57BL/6J mice

**DOI:** 10.1101/2022.04.06.487380

**Authors:** Madigan L. Bedard, Julia Sparks Lord, Patric J. Perez, Isabel M. Bravo, Adonay T. Teklezghi, Lisa Tarantino, Graham Diering, Zoe A. McElligott

## Abstract

Opioid misuse has dramatically increased over the last few decades resulting in many people suffering from opioid use disorder (OUD). The prevalence of opioid overdose has been driven by the development of new synthetic opioids, increased availability of prescription opioids, and more recently, the COVID-19 pandemic. Coinciding with increases in exposure to opioids, the United States has also observed increases in multiple Narcan (naloxone) administrations as life-saving measures for respiratory depression, and, thus, consequently, naloxone-precipitated withdrawal. Sleep dysregulation is a main symptom of OUD and opioid withdrawal syndrome, and therefore, should be a key facet of animal models of OUD. Here we examine the effect of precipitated and spontaneous morphine withdrawal on sleep behaviors in C57BL/6J mice. We find that morphine administration and withdrawal dysregulate sleep, but not equally across morphine exposure paradigms. Furthermore, many environmental triggers promote relapse to drug-seeking/taking behavior, and the stress of disrupted sleep may fall into that category. We find that sleep deprivation dysregulates sleep in mice that had previous opioid withdrawal experience. Our data suggest that the 3-day precipitated withdrawal paradigm has the most profound effects on opioid-induced sleep dysregulation and further validates the construct of this model for opioid dependence and OUD.

**Highlights:**

- Morphine withdrawal differentially dysregulates the sleep of male and female mice
- 3-day precipitated withdrawal results in larger changes than spontaneous withdrawal
- Opioid withdrawal affects responses to future sleep deprivation differently between sexes

## 1. Introduction

Opioid use disorder (OUD) is a chronically relapsing condition characterized by cycles of craving, binge, and withdrawal. In 2015, 37.8% of American adults were estimated to have used an opioid in the year prior, and from April 2020 to April 2021, opioid overdose deaths reached a record high of more than 100,000 [1,2]. Opioids have profound analgesic effects, but use may result in physical dependence, and discontinuation may lead to severe withdrawal symptoms known as opioid withdrawal syndrome (OWS), characterized by physical and affective symptoms, including weight loss, vomiting, aches, insomnia/sleep disturbances, diarrhea, irritability, dysphoria, anxiety, and social deficits [3]. In clinical populations, OWS occurs after both spontaneous and naloxone-precipitated states of withdrawal. Naloxone, an opioid receptor antagonist, is a lifesaving medication used to reverse opioid-induced respiratory depression in humans and to precipitate withdrawal in animal models of OUD [4–8]. As the availability of increasingly potent synthetic opioids enter the illicit drug market and naloxone becomes more readily available over the counter and through Emergency Medical Services (EMS), we see increased instances of patients receiving multiple administrations of naloxone, increasing 26% from 2012-2015 [9]. Recent data indicates that in Guilford County, NC, EMS administration of naloxone increased 57.8%, and repeat administration increased 84.8% compared to pre-COVID-19 pandemic numbers [10]. Furthermore, data suggests that increased education about naloxone has led to tens of thousands of overdose reversals, one such report found that 18% of clients provided naloxone reported using it to prevent overdose (for review see [11]). These data highlight the increased use of this life-saving medication, but also suggest that it is important for preclinical research to investigate how rapid precipitated withdrawal alters physiology and behavior.

With the increases in OUD and naloxone administration, it is crucial to better understand the behavioral effects of opioid withdrawal on sleep dysregulation. Sleep disruptions commonly occur during and following withdrawal from many substances, including alcohol, cannabis, and opioids [12–16]. Acute opioid administration in non-dependent adults results in sleep disturbances [17]. People with a prescription opioid dependence have been shown to differ significantly in several measures of sleep quality compared to those without dependency on prescription opioids [15]. Additionally, there is increasing evidence that sleep disruptions may be a biological pressure for relapse, especially in alcohol and opioid withdrawal [14,16,18–22]. Some of the effects of opioids on sleep have been recapitulated in preclinical models with cats, rats, and neonatal mice [23–25].

Our previous studies have utilized a model of exacerbated opioid withdrawal that leads to physical dependence. Further, these studies indicate that male and female mice experience opioid withdrawal differently, evident in their behavioral responses and neurobiology [5,26]. Far less is known about how sleep is altered during acute opioid exposure and withdrawal in mice. To our knowledge, no studies have examined the effect of opioid withdrawal on sleep behavior in female mice. One recent study highlighted how a protein that regulates circadian-dependent gene transcription altered rodent responses to fentanyl in a sex-specific manner, indicating that systems regulating circadian rhythms directly impact opioid experiences [27]. Opioid-related sleep disturbances can exacerbate both pain and withdrawal symptoms. Recent studies have shown that various opioids differentially affect sleep behaviors [28]. Zebadua Unzaga et al. (2021) found that increasing morphine and fentanyl dose-dependently affected sleep and wakefulness in C57BL/6J mice [29]. Investigating the timeline and nature of these changes and how they might differ between males and females is critical to better understand the spectrum of behavioral response, and treat opioid withdrawal. In this study, we examine how three days of repeated morphine exposure and withdrawal can acutely impact sleep behaviors and how it can alter the response to future sleep disruptions in both male and female mice.

## 2. Methods

### 2.1. Subjects

All procedures were approved by the University of North Carolina Institutional Animal Care and Use Committee. Male (N=52) and female (N=52) C57BL/6J mice (The Jackson Laboratory, Bar Harbor, ME) at least ten weeks of age were used in all experiments and were singly-housed. All animals had *ad libitum* access to food and water for the study duration. Following the conclusion of data collection, mice were euthanized according to UNC Chapel Hill’s Institutional Animal Care and Use Committee (IACUC) protocols.

### 2.2. Drugs

Morphine sulfate (Sigma-Aldrich, St. Louis, MO and Spectrum Chemical Mfg. Corp., New Brunswick, NJ) and naloxone hydrochloride dihydrate (Sigma-Aldrich, St. Louis, MO) were delivered in a 0/9% sterile saline vehicle. All injections were delivered subcutaneously at a volume of 10 mL/10kg in each mouse.

### 2.3. Piezo-Sleep System

Mice were housed individually in PiezoSleep 2.0 (Signal Solutions, Lexington, KY) 15.5 cm^2^ cages in a dedicated room separate from the general colony and behavior rooms. Mice were maintained on a 12:12 light-dark cycle (lights were set to either a 7 AM to 7 PM or a 10 am to 10 pm schedule). All data are reflected as zeitgeber time (ZT). PiezoSleep is a non-invasive housing and monitoring system that uses piezoelectric pads beneath the cage floor to detect the mouse’s movement. Using SleepStats software (SleepStats v2.18, Signal Solutions, Lexington, KY), vibrational changes were processed to determine sleeping versus waking respiratory patterns, a method validated using electroencephalography and electromyogram, and visual assessment in other studies [30–32]. Data was evaluated for several parameters including the percent time spent asleep or awake for any chosen bin size and the mean bout length of each sleep event. The first two dark cycles were discarded as habituation, and baseline data consisted of at least five days of non-disrupted behavior before any manipulations.

Here we evaluate male and female C57BL/6J mice under various conditions, including spontaneous withdrawal, precipitated withdrawal, saline control, and naloxone control (**Fig. 1**). Given our previously reported physiological and behavioral sex differences following precipitated withdrawal [5,26], and baseline sleep differences between male and female mice (**Fig. 2**), we do not statistically compare the males and females but consider qualitative comparisons. Each experiment consisted of 7 days of habituation and baseline recording, three days of withdrawal, five days of recovery recordings, a 4-hour sleep deprivation study, and additional 2 recovery days (**Fig. 1**).

**Figure 1.**
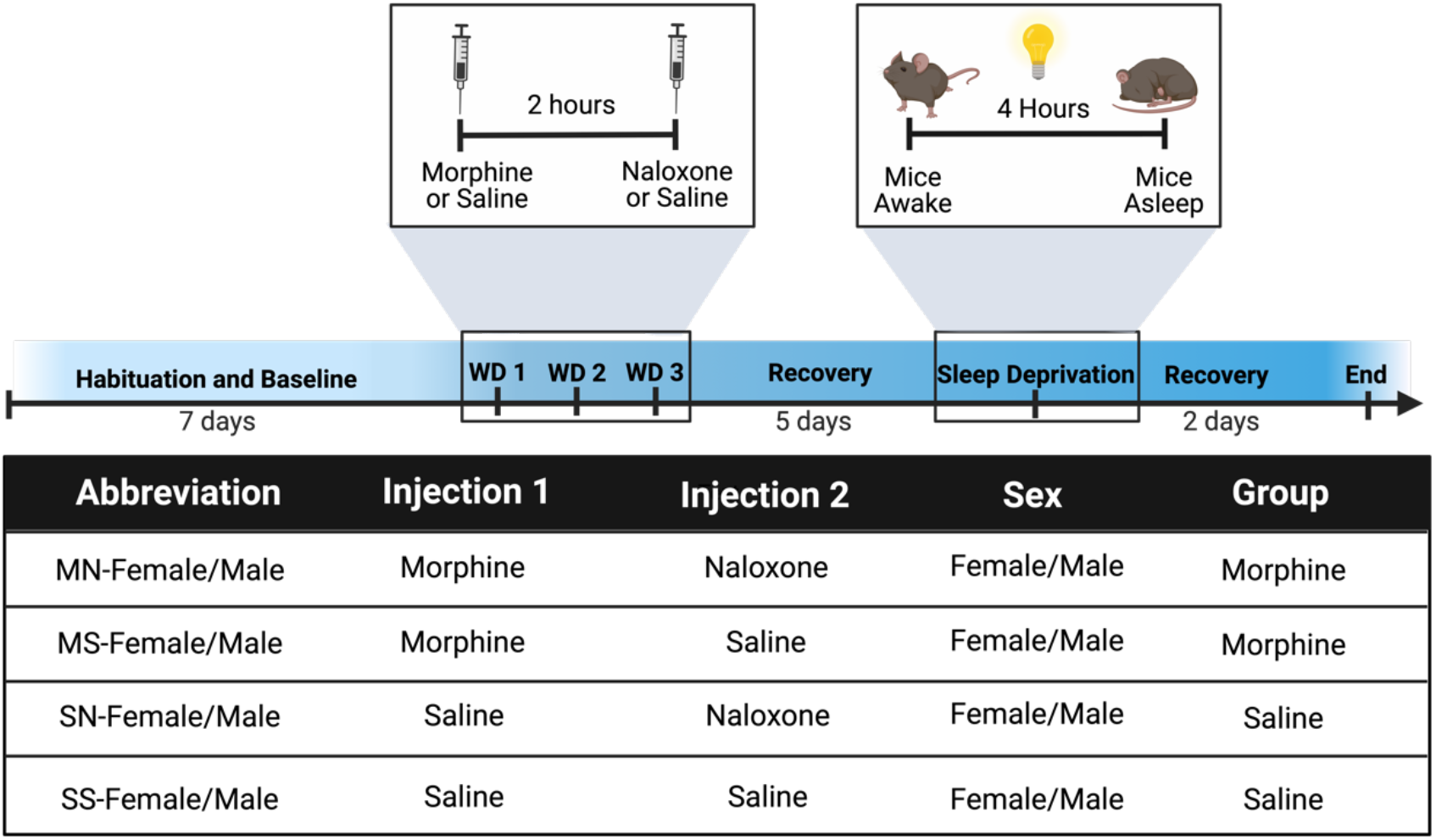
Precipitated-withdrawal paradigm in PiezoSleep chambers. Experimental timeline including habituation, baseline, morphine exposure and withdrawal, observation, and sleep deprivation. Table detailing the eight different treatment groups (four per sex).

**Figure 2.**
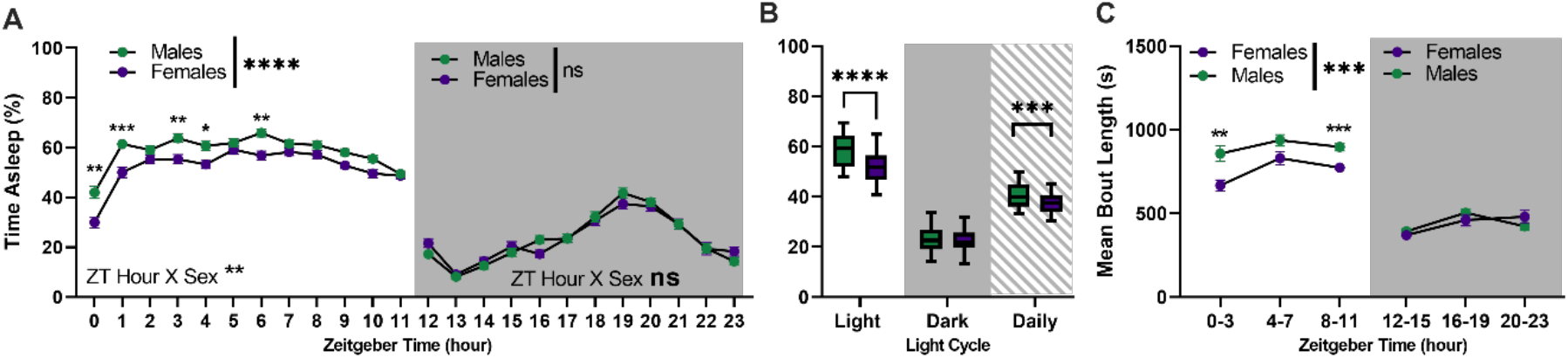
Sex differences in baseline sleep behaviors of C57BL/6J mice. Average of all baseline days shown as percent time spent sleeping of each hour in a 24-hour period, shown as Zeitgeber Time (ZT). ZT0 is the start of the light cycle and ZT12 is the start of the dark cycle. Hours 0-11 and 12-23 were analyzed separately (A). Percent time spent sleeping in a 12-hour period of all baseline days (light or dark) and across all 24-hours of all baseline days (B). Mean length (seconds) of sleep bouts in 4-hour bins across baseline days (C).

### 2.4. Withdrawal paradigm

We used a three-day withdrawal paradigm previously validated in our lab to assess the effect of morphine withdrawal on sleep behaviors in C57BL/6J mice [5,26]. This paradigm has been shown to result in sensitization of withdrawal symptoms over the three days of administration [26]. We have also shown various behavioral effects six weeks into forced abstinence using this paradigm [5]. In male Sprague-Dawley rats, a modified version of this paradigm has been shown to alter noradrenergic transmission in the ventral bed nucleus of the stria terminalis (vBNST) [33]. This paradigm allows us to investigate naloxone-precipitated withdrawal, an increasing issue in the age of synthetic opioids, and compare that state of exacerbated withdrawal with spontaneous withdrawal. Notably, we do not observe any sensitization of withdrawal signs to delivery of naloxone [26] or morphine alone (data not shown). Further, other groups using similar experimental design have highlighted the different plasticity that occurs in the brain when naloxone is used as an opioid “interrupting” agent [34,35]. Additionally, we have utilized our control groups as a way to assess the potential side effects of naloxone on the endogenous opioid system. Taken together, the four conditions help to paint a more complete picture of ceasing opioid use.

Briefly, all animals were injected with either sterile saline (0.9%) or morphine (10 mg/kg) and then 2 hours later were injected with either sterile saline (0.9%) or naloxone (1 mg/kg). Animal numbers and abbreviations for each group are as follows: male morphine naloxone (MN-Male; N=18), male saline naloxone (SN-Male; N=10), male morphine saline (MS-Male; N=16), male saline saline (SS-Male; N=8), female morphine naloxone (MN-Female; N=18), female saline naloxone (SN-Female; N=10), female morphine saline (MS-Female; N=16), female saline saline (SS-Female; N=8).

### 2.5. Sleep deprivation paradigm

All mice remained in the PiezoSleep chamber following the 3-day withdrawal paradigm. They were left untouched for five days and, on the 6^th^ day, underwent an acute sleep deprivation paradigm. Mice were kept awake during the beginning of their light cycle (ZT 0-3 for the MN-SN and MS-SS cohorts; ZT 1-4 for the MN-MS cohort). Sleep deprivation was accomplished using cage changes and gentle handling methods (light tapping on the cage top, ruffling food in the hopper, following the mouse in the cage with a pencil, removing nesting materials, etc.).

Following sleep deprivation, mice were left undisturbed for the remainder of the experiment while recordings continued.

### 2.6. Statistics

All data were analyzed using Graph Pad Prism (version 9.3.1). Data are reported as mean ± SEM. Comparisons were made with either regular 2- way ANOVAs or 3-way ANOVAs and used the Geisser-Greenhouse Correction for any post-hoc tests. All post-hoc tests were done using Bonferroni’s or Sidak’s correction. ANOVAs are reported as (F(DFn,DFd)= F-value, p= P-value). Detailed results from statistical analyses can be found in the Supplementary materials. An alpha level (p-value) of 0.05 was used for all tests. It has been established that traditional statistics do not always properly account for biological rhythms [36]. Given the expectation that there will be significant differences between light and dark cycles when assessing sleep, we conduct ANOVAs on the light cycles separately to avoid confounding biologically meaningful differences caused by our manipulations [37].

## 3. Results

### 3.1. Significant sex differences in sleep baselines

Previous studies have found differences in baseline sleep between male and female mice[38], we confirmed similar sex differences here using our sleep measurements with the PiezoSleep system. We found a main effect of sex on % time asleep in the light cycle (Figure 2A, F(1,102) =30.62, p<0.0001), and a significant interaction of sex vs. time (**Fig. 2A**, F(11,1122)=2.643, p=0.0024). Bonferroni’s multiple comparisons showed significant differences in hour 0 (p=0.0043), hour 1 (p=0.0003), hour 3 (p=0.0098), hour 4 (p=0.0398), and hour 6 (p=0.0031). We found no main effect of sex in the dark cycle (**Fig. 2A**, F(1,99) =0.0004892, p=0.9824), and no significant interaction of sex vs. time (**Fig. 2A**, F(11,1089)=1.576, p=0.1002). Bonferroni’s multiple comparisons showed no significant differences in the dark cycle.

There were no significant differences in % time asleep during the dark cycle, but males slept significantly more than females during the light cycle (**Fig. 2B**, F(1,102)=14.57, p=0.0002) and the entire day (t-test, t=3.817, df=102, p=0.0002). There was a main effect of sex on sleep bout length in the light cycle (Fig. 2C, F(1, 102) = 16.25, p=0.0001), but no significant interaction of sex vs. time in the light cycle (Fig. 2C, F(2, 204) = 1.168, p=0.3130). Bonferroni’s comparison with GG correction showed significant differences during hours 0-3 (p=0.0032) and hours 8-11 (p=0.0008). There were no main effects or statistically significant interactions in the dark cycle, though the interaction was p=0.0500 (Fig. 2C, main effect of sex: F(1, 102) = 0.01245, p=0.9114, sex x time: F(2, 204) = 3.039, p=0.0500). Because of these differences in sleep during the light cycle, and differences in sleep architecture (bouts), analyzed male and female mice separately moving forward.

### 3.2. Effect of morphine withdrawal on percent sleep in male mice

There was a main effect of the ZT hour in both ZT 0-11 and ZT 12-23 on % time asleep across all 4 days (**Supp. Table 1**). All statistics are detailed in the Supplemental and depicted within each graph as asterisks for significant effects. All groups of mice slept approximately equal amounts out of the entire day, and there were no significant differences (data not shown). Males exhibited a main effect of injection 1 (morphine vs. saline) on % time asleep across all 3 days of withdrawal and on the recovery day in both the light and dark cycles (**Fig. 3A, D, G, and J**). There was a main effect of injection 2 (naloxone vs. saline) only in the light cycle of withdrawal day 1 (WD1, **Fig. 3A**). The two injections had an interaction effect in the light cycle of WD1 (**Fig. 3A**). Injection 1 and ZT hour had significant interactions in the light cycle of the first two withdrawal days and both light cycles on the recovery day, but not the last day of withdrawal (**Fig. 3**). There were interactions between ZT hour and injection 2 in the dark cycle of WD2, light and dark of WD3, and dark cycle of the recovery day (**Fig.3 D, G, J**).

**Figure 3.**
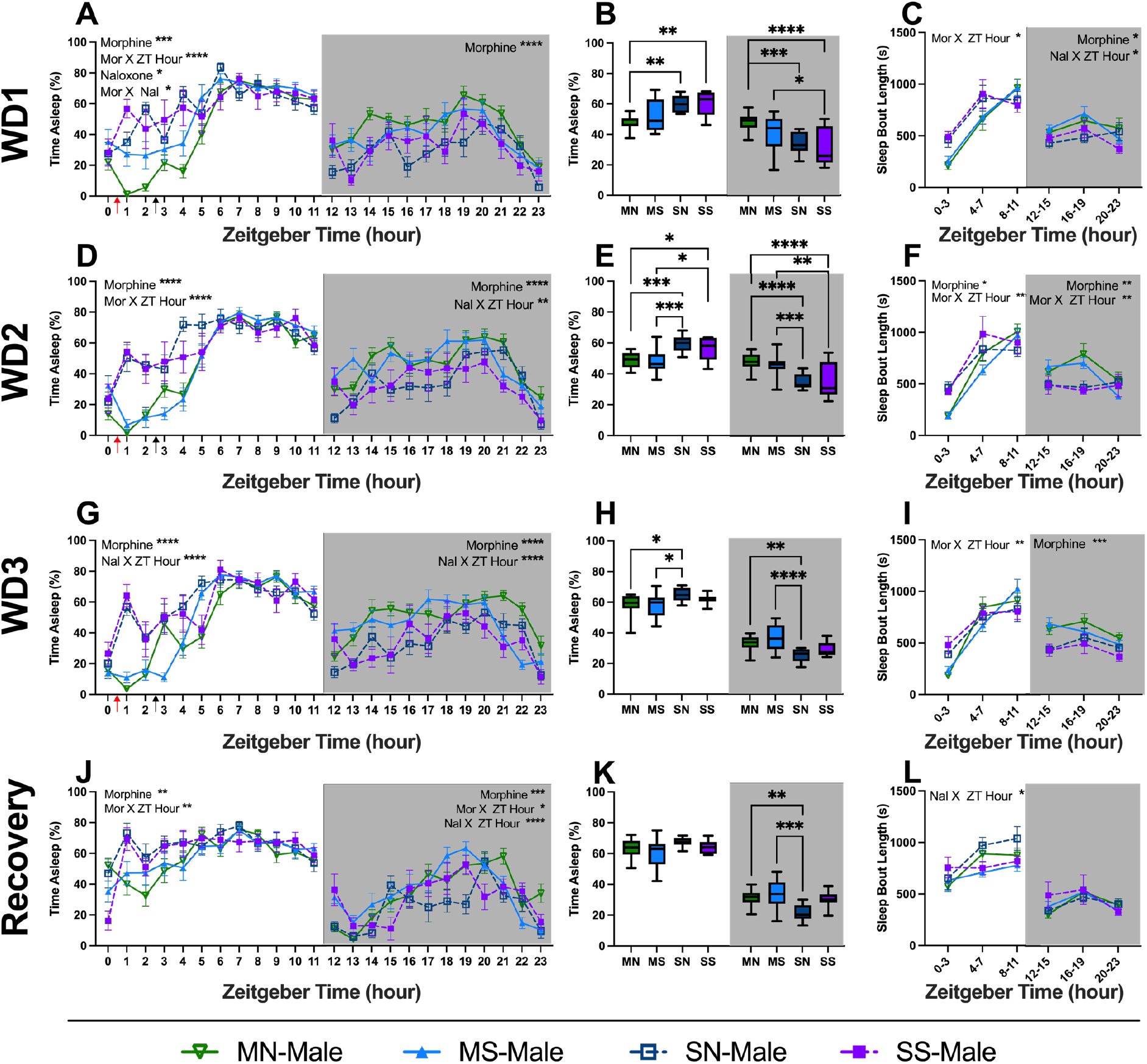
Sleep behavior of male mice during morphine exposure and withdrawal. Graphs depict either 24-hour sleep trace with percent sleep per hour, percent sleep per light cycle, or average sleep bout length in four-hour bins. Each row is a different day (withdrawal days 1-3 and then recovery day 1). Red arrows indicate morphine or saline injections (injection #1) and black arrows indicate saline or naloxone injections (injection #2). Statistical details can be found in Supplemental Tables 1 and 2. Each point or bar represents the mean ± standard error of the mean (SEM): *P < 0.05, **P < 0.01, ***P < 0.001, ****P < 0.0001. Abbreviations: Morphine-Naloxone (MN), Morphine-Saline (MS), Saline-Naloxone (SN), and Saline-Saline (SS).

Each treatment group was averaged across the light and dark cycles and assessed with a 2-way ANOVA (**Supp. Table 2**). Post-hoc tests showed significant differences within each light cycle on % time asleep during each of the three withdrawal days. On WD 1, MN-males slept less than both saline groups in the light, and MN and MS mice slept more in the dark cycle than saline controls (**Fig. 3B**; Light: MN vs. SN (p=0.002), MN vs. SS (p=0.0017) and Dark: MN vs. SN (p=0.0002), MN vs. SS (p<0.0001), and MS vs. SS (p=0.0401). On WD 2 this pattern continued and both morphine groups slept less in the light cycle and more in the dark cycle than saline groups (**Fig. 3E**; Light: MN vs. SN (p=0.0004), MN vs. SS (p=0.0497), SN vs. MS (p=0.0002), MS vs. SS (p=0.0249) and Dark: MN vs. SN (p<0.0001), MN vs. SS (p<0.0001), SN vs. MS (p=0.0005) and MS vs. SS (p=0.0014). On WD 3, morphine groups remained different than SN mice, but not SS mice (**Fig. 3H**; Light: MN vs. SN (p=0.0481), SN vs. MS (p=0.0341) and Dark: MN vs. SN (p=0.0086) and SN vs. MS (p<0.0001). Following three days of injections, the mice were left to recover on day 4 (acute withdrawal [26]) without experimental disturbance, and sleep behavior was continuously monitored. Males exhibited lasting changes into the dark cycle of the recovery day (**Fig. 3K**; Dark: MN vs. SN (p=0.0028) and MS vs. SN (p=0.0001)).

Sleep bout lengths also differed between the groups. There was an interaction between injection 1 and ZT hour in the light cycle (**Fig. 3C**; Light: F (2, 96) = 6.089, p=0.0032). In the dark cycle, morphine significantly increased sleep bout lengths (**Fig. 3C**; Dark: F (1, 48) = 5.521, p=0.0229) and naloxone had a significant interaction with ZT hour (**Fig. 3C**; Dark: F (2, 96) = 3.831, p=0.0251). On WD 2, morphine altered sleep bout lengths (**Fig. 3F**; Light: F (2, 96) = 6.089, p=0.0032; Dark: F (1, 48) = 8.457, p=0.0055) and an interaction of injection 1 with ZT hour (Light: F (2, 96) = 9.242, p=0.0002; Dark: F (2, 96) = 5.932, p=0.0037). By WD 3, there was a main effect of injection 1 on sleep bout lengths, but only in the dark (**Fig. 3I**; Dark: F (1, 48) = 13.54, p=0.0006) and an interaction of injection 1 with ZT hour in the light (Light: F (2, 96) = 6.753, p=0.0018). Finally, there was an interaction between injection 2 and ZT hour in the light cycle on the recovery day (**Fig. 3L**; Light: F (2, 96) = 4.620, p=0.0121).

### 3.3. Effect of morphine withdrawal on percent sleep in female mice

As with the males, there was a main effect of the ZT hour in both ZT 0-11 and ZT 12-23 on % time asleep across all 4 days (**Supp. Table 3**). All statistics are detailed in the supplemental table and depicted within each graph as asterisks for significant effects. All groups of mice slept approximately equal amounts of the entire day, and there were no significant differences (data not shown).

Females also exhibited a main effect of injection 1 (morphine vs. saline) across all 3 days of withdrawal and on the recovery day in both the light and dark cycles (**Fig. 4A, D, G, and J**). There was no main effect of injection 2 (naloxone vs. saline). Injection 1 and ZT hour had significant interactions in the light cycle of all 4 days and the dark cycles on WD 2 and the recovery day (**Fig. 4**). There were interactions between ZT hour and injection 2 in the dark cycle of all 4 days and the light cycle of WD 1 (**Fig. 4A, D, G, and J**).

**Figure 4.**
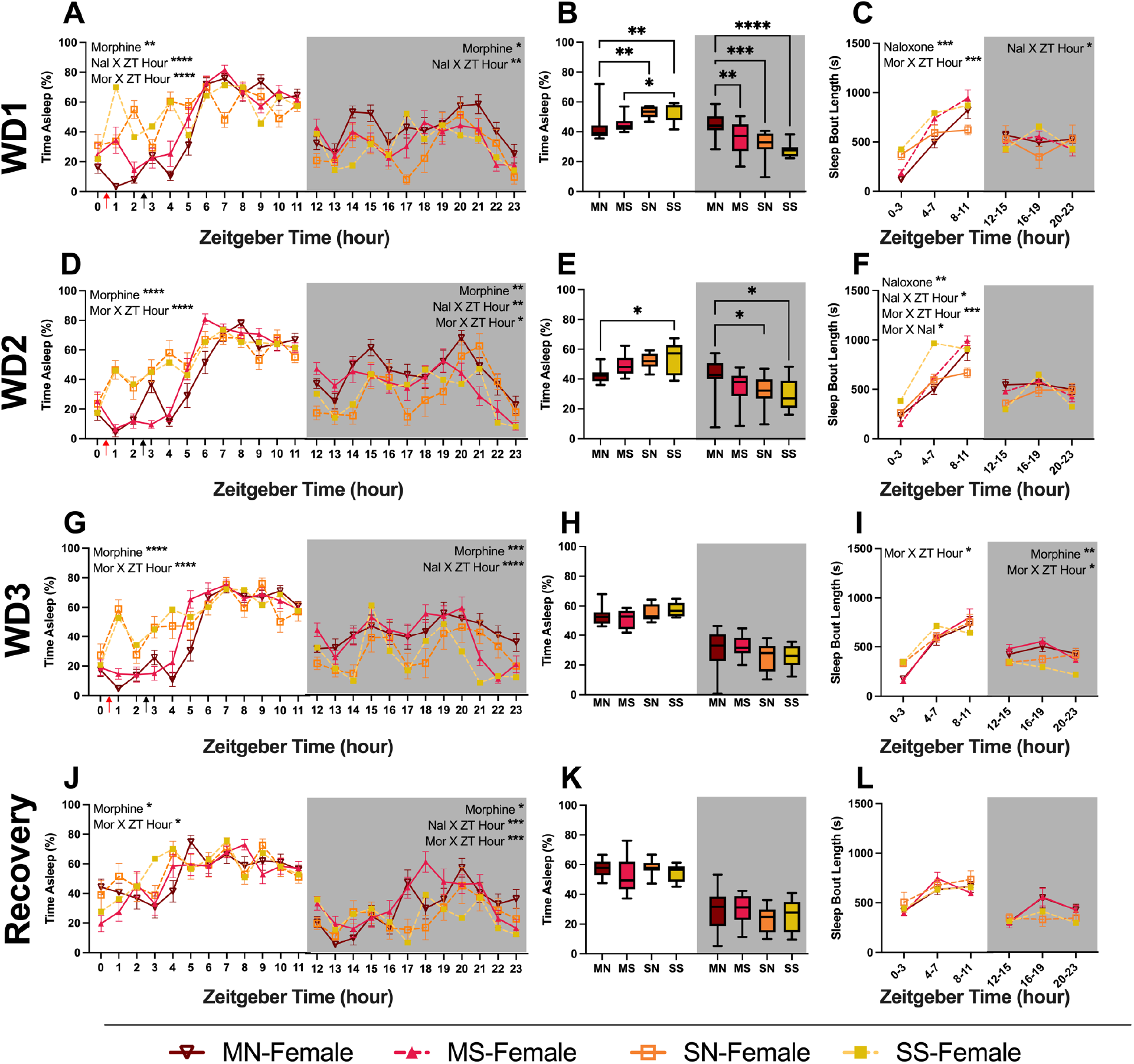
Sleep behavior of female mice during morphine exposure and withdrawal. Graphs depict either 24-hour sleep trace with percent sleep per hour, percent sleep per light cycle, or average sleep bout length in four-hour bins. Each row is a different day (withdrawal days 1-3 and then recovery day 1). Red arrows indicate morphine or saline injections (injection #1) and black arrows indicate saline or naloxone injections (injection #2). Statistical details can be found in Supplemental Tables 2 and 3. Each point or bar represents the mean ± standard error of the mean (SEM): *P < 0.05, **P < 0.01, ***P < 0.001, ****P < 0.0001. Abbreviations: Morphine-Naloxone (MN), Morphine-Saline (MS), Saline-Naloxone (SN), and Saline-Saline (SS).

Each treatment group was averaged across the light and dark cycles and 2-way ANOVAs were run (**Supp. Table 2**). Post-hoc tests showed significant differences only on the first two withdrawal days. On WD 1, morphine females slept less than both saline mice in the light, and more in the dark cycle than the other three groups (**Fig. 4B**; Light: MN vs. SN (p=0.004), MN vs. SS (p=0.004), MS vs. SS (p=0.0455) and Dark: MN vs. SN (p=0.0004), MN vs. MS (p=0.0091), and MN vs. SS (p<0.0001). On WD 2 this pattern weakened and only the MN slept less in the light cycle than the saline controls (**Fig. 4E**; Light: MN vs. SS (p=0.0207). In the dark cycle, MN mice slept more than both saline groups (**Fig. 4E**; Dark: MN vs. SN (p=0.042) and MN vs. SS (p=0.016)).

Sleep bout lengths also differed between the groups. On WD 1, there was a main effect of naloxone on sleep bout length in the light cycle (**Fig. 4C**; Light: F (1, 48) = 14.41, p=0.0004). There was an interaction between injection 1 and ZT hour in the light cycle (**Fig. 4C**; Light: F (2, 96) = 8.255, p=0.0005). In the dark cycle, injection 2 had a significant interaction with ZT hour (**Fig. 4C**; Dark: F (2, 96) = 4.627, p=0.0121. On WD 2, naloxone altered sleep bout lengths (**Fig. 4F**; Light: F (1, 48) = 11.48, p=0.0014). There were also multiple interactions in the light cycle (**Fig. 4F**; Inj 1 X ZT hour: F (2, 96) = 9.005, p=0.0003; Inj 2 X ZT hour: F (2, 96) = 3.190, p=0.0456; Inj 1 X Inj 2: F (1, 48) = 5.379, p=0.0247). On WD 3, there was an interaction between injection 1 and ZT hour in both light cycles (**Fig. 4I**; Light: F (2, 96) = 3.128, p=0.0483); Dark: F (2, 96) = 3.606, p=0.0309) and morphine resulted in longer sleep bouts in the dark cycle (F (1, 48) = 12.02, p=0.0011).

### 3.4. Cumulative sleep displaced from baseline following withdrawal

To investigate how individual animals shifted their sleep patterns during the treatment and withdrawal from opioids, we also examined the data as cumulative minutes of sleep displaced from baseline. Percentages were converted to total minutes spent asleep and then subtracted from the total minutes slept up to that point during baseline for each individual animal [39]. This method of analysis allows for easier between-group assessments while also normalizing each mouse to itself. We then fit linear regression models to each line, shown on separate graphs for ease of visualization. Additionally, we compare the cumulative difference from baseline at hour 23 (total minutes different from baseline across an entire day).

There are significant effects of morphine, significant interactions of naloxone X ZT hour, and significant interactions of morphine X ZT hour across all 4 days in both males (**Fig. 5A, D, G, J; Supp. Table 4**) and females (**Fig. 6A, D, G, J; Supp. Table 5**). Naloxone significantly impacted males on WD1, WD3, and recovery (**Fig. 5A, G, J; Supp. Table 4**), but it only altered sleep in females during the withdrawal days, not recovery (**Fig. 7A, D, G; Supp. Table 5**).

**Figure 5.**
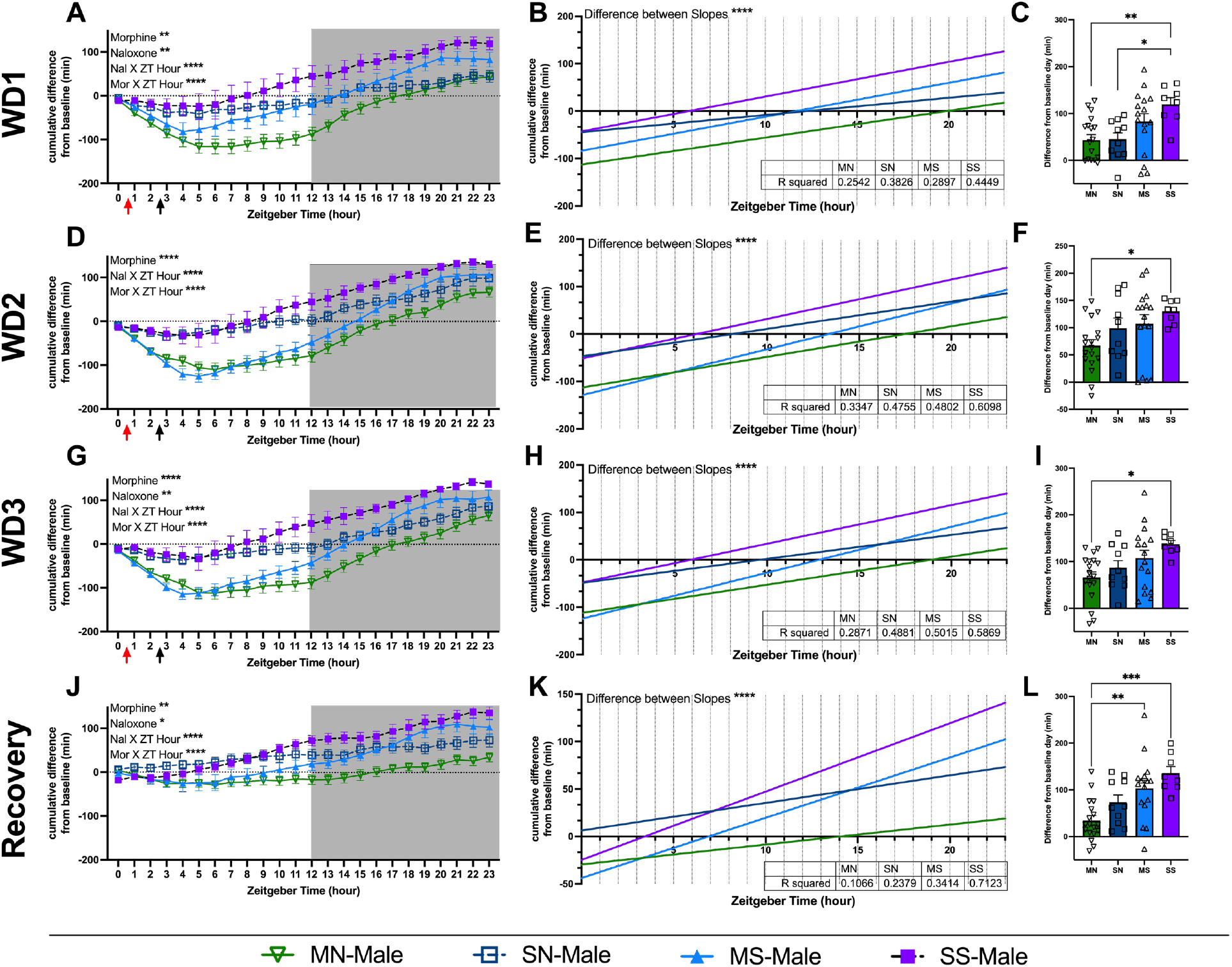
Cumulative difference in minutes of sleep of male mice during morphine exposure and withdrawal. Minutes of sleep compared to each animal’s baseline sleep (within subjects). (**A,D, G, and J**) Graphs show minutes of sleep subtracted from baseline minutes, cumulative by hour. Red arrows indicate morphine or saline injections (injection #1) and black arrows indicate saline or naloxone injections (injection #2). Each row is a different day (withdrawal days 1-3 and then recovery day 1). Grey shading shows the dark cycle. Data above 0 indicate a mouse had slept more by that point in the day than they did by that same hour during their baseline. (**B, E, H, and K**) Graphs show linear regression of within-animal graphs. R-squared values are reported in the inset tables. (**C, F, I, and L**) Graphs show the final difference in minutes slept over 24 hours compared to baseline. Statistical details can be found in Supplemental Table 4. Each point represents the mean ± standard error of the mean (SEM): *P < 0.05, **P < 0.01, ***P < 0.001, ****P < 0.0001. Abbreviations: Morphine-Naloxone (MN), Morphine-Saline (MS), Saline-Naloxone (SN), and Saline-Saline (SS).

**Figure 6.**
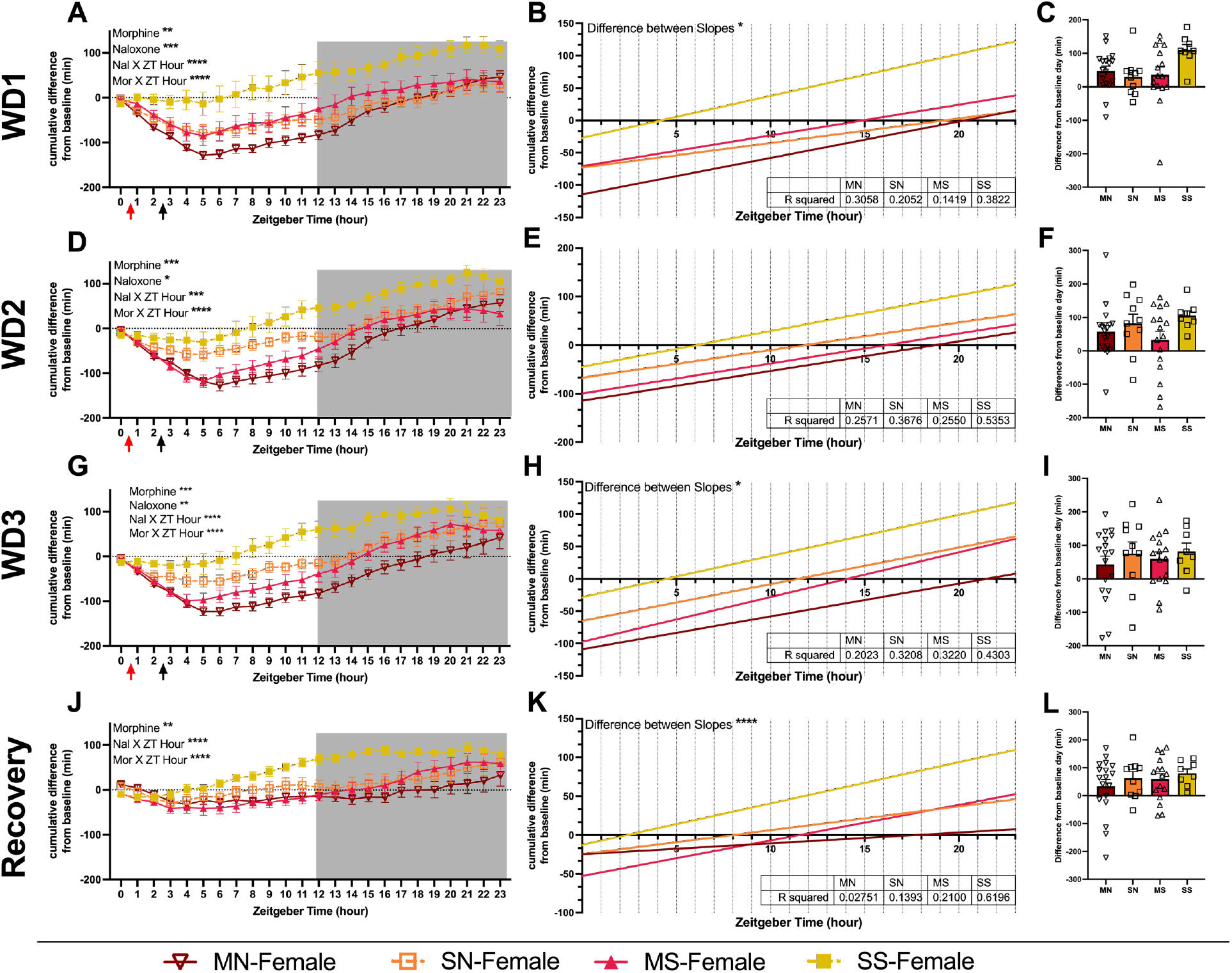
Cumulative difference in minutes of sleep of female mice during morphine exposure and withdrawal. Minutes of sleep compared to each animal’s baseline sleep (within subjects). (**A,D, G, and J**) Graphs show minutes of sleep subtracted from baseline minutes, cumulative by hour. Red arrows indicate morphine or saline injections (injection #1) and black arrows indicate saline or naloxone injections (injection #2). Each row is a different day (withdrawal days 1-3 and then recovery day 1). Grey shading shows the dark cycle. Data above 0 indicate a mouse had slept more by that point in the day than they did by that same hour during their baseline. (**B, E, H, and K**) Graphs show linear regression of within-animal graphs. R-squared values are reported in the inset tables. (**C, F, I, and L**) Graphs show the final difference in minutes slept over 24 hours compared to baseline. Statistical details can be found in Supplemental Table 5. Each point represents the mean ± standard error of the mean (SEM): *P < 0.05, **P < 0.01, ***P < 0.001, ****P < 0.0001. Abbreviations: Morphine-Naloxone (MN), Morphine-Saline (MS), Saline-Naloxone (SN), and Saline-Saline (SS).

**Figure 7.**
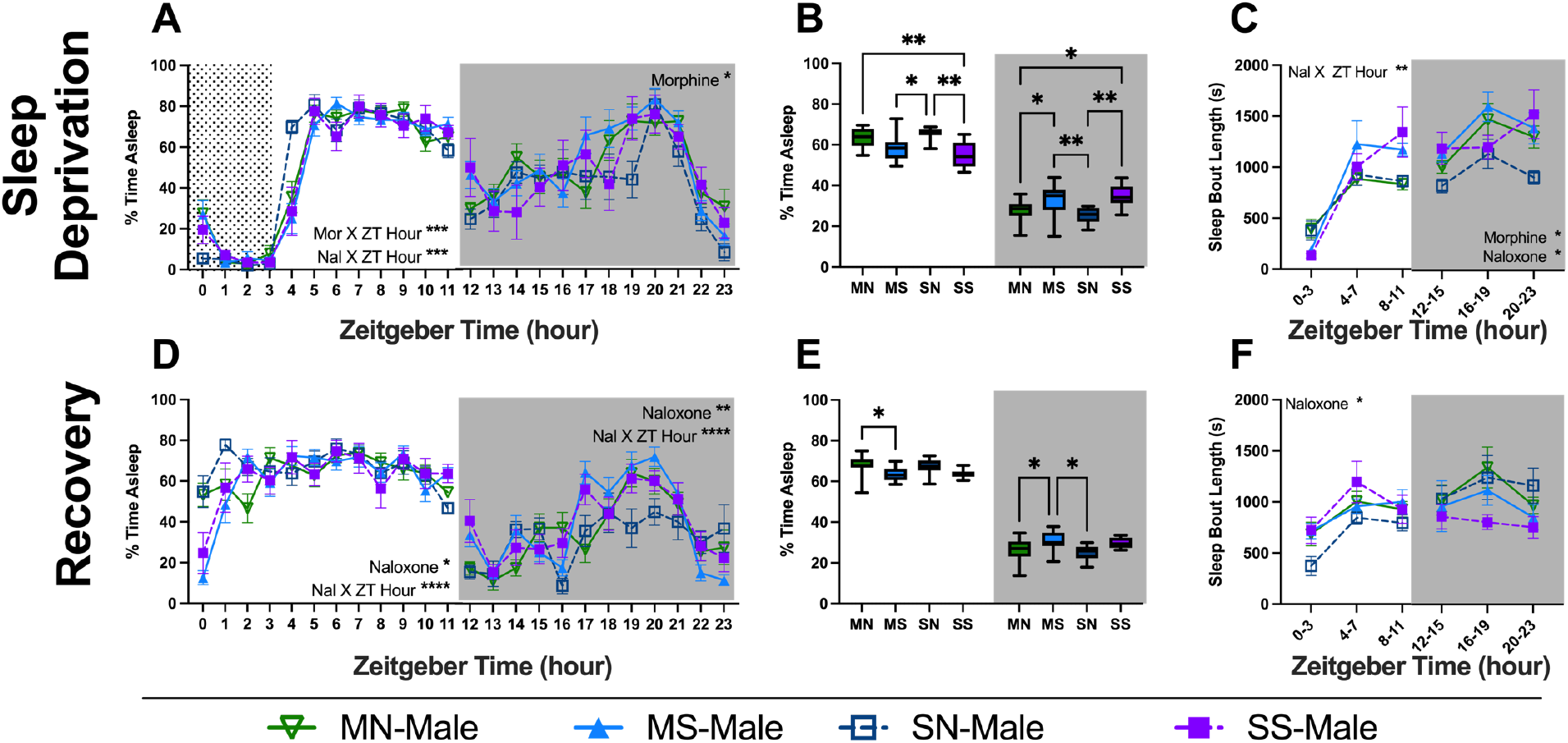
Sleep behavior of male mice during sleep deprivation and recovery. Sleep percentage of male mice undergoing sleep deprivation for 4 hours (**A-C**) and the day immediately following (**D-L**). Graphs are either 24-hour sleep percentages (**A, D**), sleep percentages by light cycle (**B, E**), or sleep bout length in 4-hour bins (**C,F**). Grey box shows lights off and dotted region shows sleep disruption period. Statistical details can be found in Supplemental Table 6. Each point or bar represents the mean ± standard error of the mean (SEM): *P < 0.05, **P < 0.01, ***P < 0.001, ****P < 0.0001. Abbreviations: Morphine-Naloxone (MN), Morphine-Saline (MS), Saline-Naloxone (SN), and Saline-Saline (SS).

We also performed a linear regression on the within-data graphs, resulting in the regression lines seen in figures 5 and 6 (**Fig. 5-6: B, E, H, K; Supp. Tables 4 & 5**). In males and females, all lines were significantly different from zero, which indicates the mice all slept differently from baseline conditions. The slopes of each line were significantly different across all days for males (**Fig. 5**), but only on some of the days for females (**Fig. 6**).

Finally, we evaluated the total difference from baseline between each group in the last column of figures 5 and 6. All mice across treatment groups and sexes slept more than they did at baseline. This indicates a natural increase in sleep in response to handling and injections.

Indeed, we were very surprised how drastically the SS mice showed differences in sleep behavior as compared to their baseline even on the recovery day when they were not perturbed. Males who received naloxone slept most similarly to their baseline times on WD1, and slept significantly less than SS mice (**Fig. 5C;** MN vs SS p=0.0081, SN vs. SS p=0.0251). MN-males also differed from SS on WD2 (**Fig. 5F**; p=0.0422), WD3 (**Fig. 5I**; p=0.0134), and recovery day (**Fig. 5L**; p=0.0002). MN-males also slept less than MS-males by the recovery day (**Fig. 5L**; p=0.0024). Female mice showed no differences between groups in how much they differed from their baseline (**Fig. 6**). Female mice had much greater within group variation than the male mice.

### 3.5. Sleep deprivation in male and female mice

Following 6 days of forced abstinence where the mice were not manipulated by the investigators, we next conducted sleep deprivation assays. All mice were kept awake for 4 hours at the beginning of their light cycle, staring at either ZT0 or ZT1 (see methods). On the sleep deprivation day, males showed significant interactions of each injection with ZT hour in the light cycle and main effect of the morphine injection in the dark cycle (**Fig. 7A; Supp. Table 6**). Females showed similar interactions in the light cycle but also had an interaction of naloxone and ZT hour (**Fig. 8A; Supp. Table 7**). Male mice exhibited more pronounced effects on the recovery day than the females with significant interactions between naloxone and ZT hour as well as a main effect of naloxone in both light cycles, whereas females only showed a naloxone X ZT hour interaction in the dark cycle (**Fig. 7D & 8D**). Males also showed differences within each light cycle between groups on both the sleep deprivation and recovery day. In general, the mice who received naloxone slept differently than those who did not, regardless of morphine exposure on the sleep deprivation day (**Fig. 7B**). Also, both injections altered sleep bout length in the dark cycle (**Fig. 7C**). On the recovery day, naloxone-receiving mice slept more in the light and less in the dark cycle, and had altered bout lengths compared to mice who did not get naloxone (**Fig. 7E & F**). Female mice showed no differences between groups when averaged across 12 hours or when looking at bout length (**Fig. 8**).

**Figure 8.**
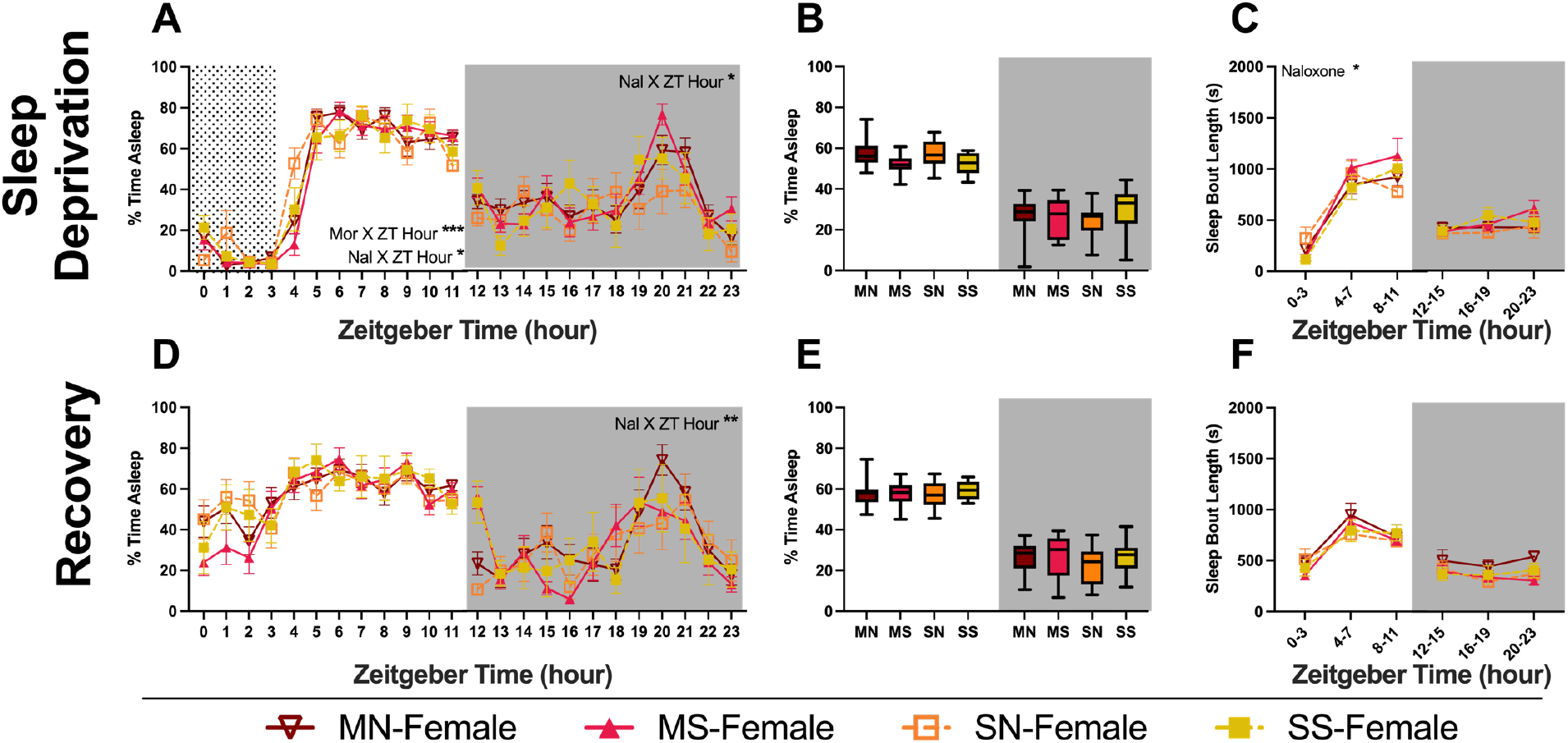
Sleep behavior of female mice during sleep deprivation and recovery. Sleep percentage of male mice undergoing sleep deprivation for 4 hours (**A-C**) and the day immediately following (**D-L**). Graphs are either 24-hour sleep percentages (**A, D**), sleep percentages by light cycle (**B, E**), or sleep bout length in 4-hour bins (**C,F**). Grey box shows lights off and dotted region shows sleep disruption period. Statistical details can be found in Supplemental Table 7. Each point or bar represents the mean ± standard error of the mean (SEM): *P < 0.05, **P < 0.01, ***P < 0.001, ****P < 0.0001. Abbreviations: Morphine-Naloxone (MN), Morphine-Saline (MS), Saline-Naloxone (SN), and Saline-Saline (SS).

### 3.6. Cumulative minutes displaced from baseline sleep during sleep deprivation and recovery

As with the withdrawal days, we normalized each mouse’s sleep to their own baseline (prior to any manipulations) and then compared between groups (**Supp. Table 8**). This revealed main effects of naloxone in males (**Fig. 9A**) and morphine in females (**Fig. 10A**) on the sleep deprivation day. Both sexes showed significant interactions on both sleep deprivation and recovery days (**Fig. 9A, D & 10A, D**). Both sexes were fit with linear regressions and the difference between the slopes was significant in all cases (**Fig. 9 B, E & 10B, E**). All male groups slept more total time on sleep deprivation and recovery than they did on baseline days (**Fig. 9 C,F**). However, the increase in MN-males was significantly less than SS-males on both days (**Fig. 9 C,F**). Female mice showed no overall differences between groups in their increase from baseline (Fig. 10 C,F). All groups in both sexes slept more than their baselines on average, but individual variability is clear with some animals sleeping less than they did at baseline (**Fig. 9 & 10**).

**Figure 9.**
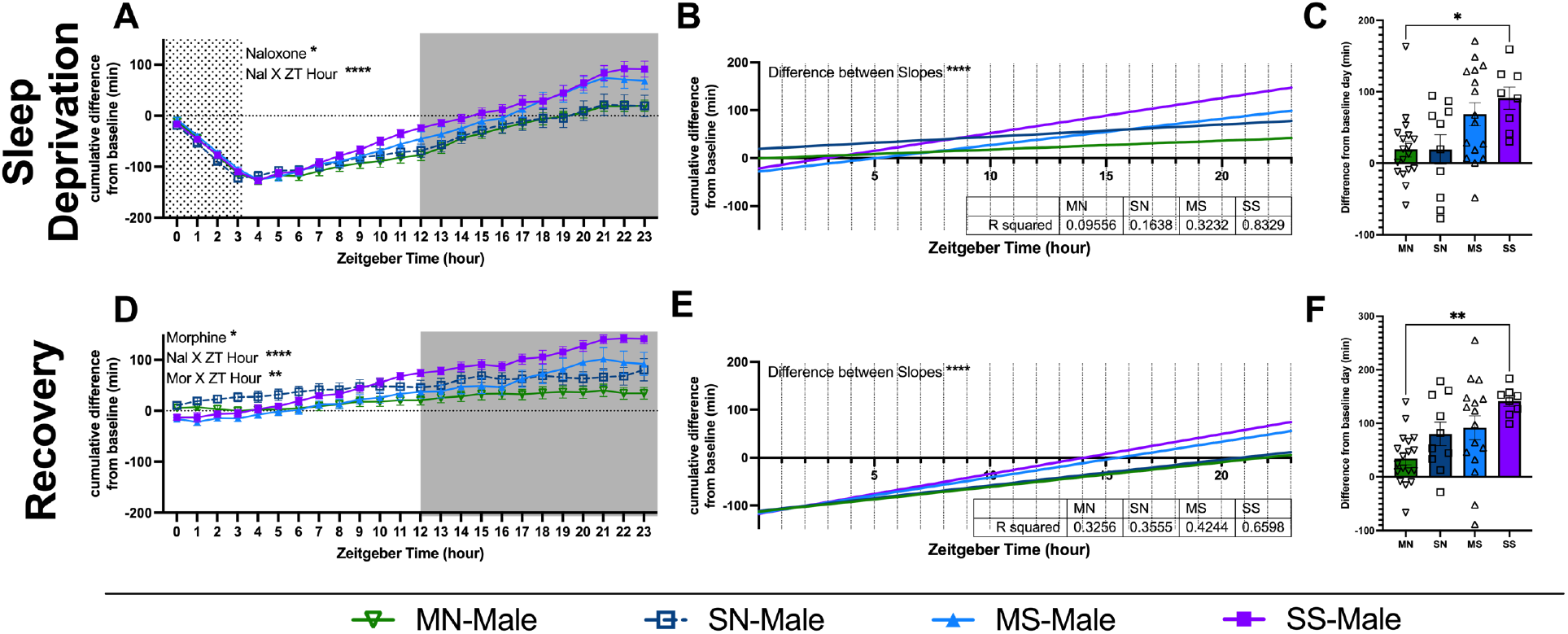
Cumulative difference in minutes of sleep of male mice during sleep disruption and recovery. Minutes of sleep compared to each animal’s baseline sleep (within subjects). (**A, D**) Graphs show minutes of sleep subtracted from baseline minutes, cumulative by hour. Each row is a different day (sleep deprivation and recovery day). Grey shading shows the dark cycle. Data above 0 indicate a mouse had slept more by that point in the day than they did by that same hour during their baseline. (**B, E**) Graphs show linear regression of within-animal graphs. R-squared values are reported in the inset tables. (**C, F**) Graphs show the final difference in minutes slept over 24 hours compared to baseline. Statistical details can be found in Supplemental Table 8. Each point represents the mean ± standard error of the mean (SEM): *P < 0.05, **P < 0.01, ***P < 0.001, ****P < 0.0001. Abbreviations: Morphine-Naloxone (MN), Morphine-Saline (MS), Saline-Naloxone (SN), and Saline-Saline (SS).

**Figure 10.**
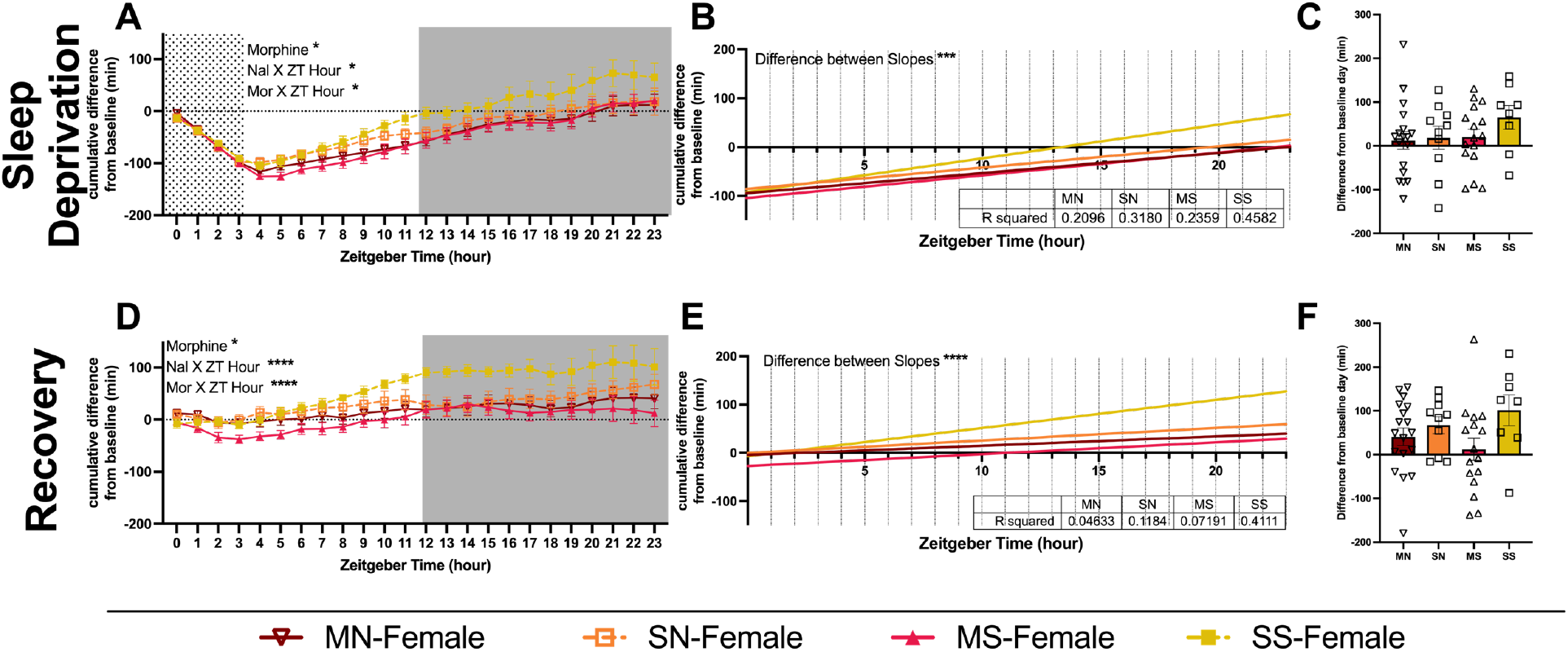
Cumulative difference in minutes of sleep of female mice during sleep deprivation and recovery. Minutes of sleep compared to each animal’s baseline sleep (within subjects). (**A, D**) Graphs show minutes of sleep subtracted from baseline minutes, cumulative by hour. Each row is a different day (sleep deprivation and recovery day). Grey shading shows the dark cycle. Data above 0 indicate a mouse had slept more by that point in the day than they did by that same hour during their baseline. (**B, E**) Graphs show linear regression of within-animal graphs. R-squared values are reported in the inset tables. (**C, F**) Graphs show the final difference in minutes slept over 24 hours compared to baseline. Statistical details can be found in Supplemental Table 8. Each point represents the mean ± standard error of the mean (SEM): *P < 0.05, **P < 0.01, ***P < 0.001, ****P < 0.0001. Abbreviations: Morphine-Naloxone (MN), Morphine-Saline (MS), Saline-Naloxone (SN), and Saline-Saline (SS).

## 4. Discussion

Sleep is governed by multiple brain regions and neurotransmitters. EEG studies have identified the various regions involved in stages of sleep including portions of cortex, thalamus, hypothalamus, hippocampus, basal forebrain and others [40–42]. Many brain regions known to be involved in sleep are also implicated in addiction/reward circuitry and stress systems. In particular, monoaminergic nuclei have been largely implicated in regulating sleep behavior. The locus coeruleus (LC) projects to the basal forebrain, bringing dense noradrenergic inputs that are known to correlate with arousal states. The LC is the beginning of a wake-promoting circuit that uses monoaminergic signaling through the midbrain and to the frontal cortex and receives feedback from orexinergic neurons in the lateral hypothalamus [44]. Increased activation of the LC promotes arousal, and decreased activation of the LC results in increases in NREM sleep [44]. During opioid withdrawal, the LC neurons are activated and there is an increase in norepinephrine in the ventral forebrain that drives opioid withdrawal-induced aversion [45].

There is also an increase in norepinephrine in the ventral bed nucleus of the stria terminalis (vBNST) following opioid withdrawal, although the source of this NE is mainly from the medullary noradrenergic neurons which have a less well-defined role in sleep [46,47]. The ventral tegmental area (VTA) is considered the preeminent common substrate for all rewarding substances and reinforcing behaviors. It sends dopamine to the nucleus accumbens and is highly implicated in facilitating rewarding behaviors and self-administration of drugs. The VTA consists of both dopaminergic and non-dopaminergic (GABAergic) neurons, each of which have been shown to play a role in the control of sleep behaviors [48,49]. The GABAergic neurons in the VTA show increased firing during REM sleep and decreased firing during waking behaviors [48]. These neurons synapse onto other GABA-neurons as well as onto the DA neurons in the VTA [50]. Inhibition of these dopamine neurons results in sleep induction and sleep preparation behaviors such as nest-building [49].

Here we examined how spontaneous and precipitated opioid withdrawal modulates sleep behavior in male and female mice using a non-invasive sleep measurement across multiple days and during a 4-hour sleep deprivation and recovery. We compare a 3-day precipitated morphine withdrawal paradigm that we and others have used to model the development of opioid dependence/tolerance as compared to spontaneous withdrawal from the same dose of morphine. We also compare these paradigms to their respective controls (SS and SN) measured concurrently. While no animal model can completely capture the complexity of the human condition, this model captures elements in the following ways: (1) Upwards of 25% of patients receiving opioid therapies in a primary care setting can develop opioid use disorder [51,52], therefore it is critical to model how withdrawal from relatively low doses of opioids (here we use a 10 mg/kg dose in the mouse, an analgesic dose for male C57BL/6J mice). (2) There is increased use of naloxone as a critical life saving measure to reverse overdose and there is evidence of repeated administrations in clinical populations, ergo we need to have an understanding of how this treatment affects brain circuitry and behavior [9,10]. (3) We observe behavioral and plastic changes both in an “acute” withdrawal phase (24 hours following withdrawal) and in a protracted phase (6 weeks into forced abstinence) demonstrating the lasting effects of this paradigm on neural function [5,26]. These adaptations are akin to those observed by other researchers, notably the Rothwell lab [34,35].

Generally, our data suggest that repeated withdrawal dysregulates sleep behavior, and the precipitated withdrawal paradigm results in greater sleep dysregulation, including subsequent drug-free recovery days and alterations in sleep drive following sleep deprivation. Additionally, we observed differences in some of these metrics between male and female subjects, and confirm sleep differences at baseline between male and female C57BL6/J mice.

### 4.1. Difference in baseline sleep behaviors in male and female mice

In this study, we compared our model of OUD – a 3-day precipitated morphine withdrawal paradigm which results in acute withdrawal responses (ie. sensitization of escape jumps, paw tremors, and fecal boli over 3 days [26]) as well as protracted withdrawal responses (ie. changes in open field behaviors, social interaction, and increased locomotion[5]) -- to spontaneous morphine withdrawal, in the context of sleep dysregulation. Our previous studies also also uncovered sex differences that occur during both acute and protracted withdrawal. We conducted these paradigms inside our PiezoSleep chambers and examined the effects of morphine withdrawal on the sleep behaviors of male and female C57BL/6J mice (**Fig. 1**). Similar to other reports in rodents[38], we saw that male and female C57BL/6J mice sleep differently at baseline, with the males sleeping on average 58.3% of the light cycle and 22.9% of the dark cycle while females slept 52.0% of the light cycle and 22.6% of the dark cycle (**Fig. 2**).

Additionally, the mean sleep bout length varied between male and female mice (**Fig. 2**). This pattern in sleep differences between males and females is consistent with observations in healthy adult humans [53]. Once we established baseline sleep behaviors, mice were exposed to morphine followed by withdrawal. Due to the significant difference between male and female baselines, the sexes are only compared qualitatively for the remainder of the discussion.

### 4.2. Validation of effects during precipitated withdrawal paradigm

We showed that our naloxone-precipitated morphine withdrawal paradigm results in sleep disruptions similar to observations in clinical reports from patients with OUD [14,15,22]. Morphine results in decreased sleep in the light cycle and increased sleep in the dark cycle for both males and females across all three days of injections (**Fig. 3 & 4**). We observed correlating changes in the length of sleep bouts. Morphine exposure resulted in decreased bout length in the light cycle and increased bout length in the dark cycle (**Fig. 3 & 4**). This parallels the human experience in which those taking opioids or withdrawn from opioids report increased daytime sleepiness followed by insomnia at night [54–57]. Naloxone injections, on the other hand, result in an immediate increase in sleep in either male or female animals (**Fig. 3 & 4: A, D, G**). In non-opioid-dependent people, naloxone alone results in increased latency to reach REM sleep, duration of REM, and the number of REM cycles [58]. These effects are likely due to activation of the LC, which only ceases firing during REM sleep. In those with OUD, maintenance treatment with buprenorphine and naloxone (Suboxone) results in improved sleep compared to those going through treatment without the combination therapy, but the sleep patterns do not return to baseline [59–61]. Across all of our mice, the total daily percent time spent sleeping and daily mean bout length were not different between groups on any given experimental day (data not shown). This result suggests that sleep displaced by morphine exposure and withdrawal was regained at other time periods during the day. It is unlikely that this finding is consistent with human behavior given the environmental and social pressures to be active or productive during the day. For example, one study found that illicit opioid use decreased the amount of sleep its participants achieved, but so did having earlier scheduled urinalysis appointments [62]. These data highlight that the effects of sleep alterations could be more impactful in humans. Future studies could mimic the requirements/daily pressures of human life, such as working for food or sleep deprivation (see below), and explore how these changes further impact sleep behavior.

### 4.3. Sleep disruptions continue into the first day of abstinence in both male and female mice

While the 3-day precipitated withdrawal paradigm was originally developed to promote the rapid development of opioid dependence [63], the current opioid epidemic has resulted in people receiving doses of Narcan® (naloxone) to alleviate respiratory depression and save lives. The rates of people receiving multiple doses of Narcan from Emergency Medical Services is on the rise (up 26% from 2012-2015 [9] and up 84.8% in Guilford County, NC from pre-COVID to 2021 [10]). The increasing frequency of repeated withdrawals and repeated dosing of naloxone begs the question of what rapid withdrawal itself, not merely opioid exposure and spontaneous withdrawal, does to the brain. Here, we see that there are acute effects of morphine withdrawal on sleep that persist into the day following the last exposure and withdrawal. Both males and females show interactions between the hour and treatment group into abstinence (**Fig. 3,4: J**). MN-male and MS-male mice showed a significant reduction in sleep during the light phase (similar to the time point where they received morphine/naloxone on the preceding days), as well as an increase in sleep during the dark when compared to their saline-naloxone controls (**Fig. 3K**). This is not the case, however, for females (**Fig. 4K**). The persistence of disruptions 24-36 hours after the last dose of morphine is comparable to that seen in the human condition, where studies have shown disruptions may never completely disappear due to external stressors (see discussion on sleep deprivation below) [60,61]. At this same 24-hour time point, our lab has shown that GABAergic signaling in the BNST is altered following opioid withdrawal [26]. Intriguingly, the plasticity was different in males and females: males exhibited increased frequency of spontaneous IPSCs, while the females showed decreased frequency of sIPSCs.

An increase in GABAergic signaling in the BNST might result in a disinhibition of VTA GABA neurons, which have been shown to have decreased activity during awake states [26,48]. This circuit provides a potential explanation given the differences in GABA signaling and sleep behaviors in acute withdrawal, and warrants further investigation. We have also examined various behaviors 6 weeks following precipitated withdrawal and seen several sex differences [5]. These data indicate that in the future, it might be beneficial to observe sleep behavior at a more protracted time point similar to our previous studies of protracted withdrawal behavior [5].

### 4.4. Within-animal comparisons elucidate differential responses by male and female mice

In addition to assessing the sleep percentage by hour, we have examined sleep behavior by calculating the cumulative difference in minutes of sleep obtained throughout the day compared to baseline sleep. This form of analysis considers within-subject variability and allows each mouse to serve as their own baseline. Here, we can more clearly visualize the effect of naloxone in combination with morphine on male mice (**Fig. 5**). On the recovery day, males, but not females, exhibited a significant main effect of both morphine and naloxone (**Fig. 5 & 6: J**). This finding is particularly interesting given that precipitated and spontaneous withdrawal groups behaved almost identically to each other on the three days of withdrawal (**Fig. 5C, F, I**).

However, during acute withdrawal, precipitated withdrawal animals slept significantly less than spontaneous withdrawal animals, indicating a role of withdrawal severity on symptoms (**Fig. 5L**). For the females, spontaneous and precipitated withdrawal groups behaved much more similarly, indicated by only a main effect of morphine but not naloxone (**Fig. 6**). These findings indicate that male mice may be more affected by the combination of morphine and naloxone than female mice, specifically at the 1mg/kg dose of naloxone. One clinical study showed that women are more likely than men to use opioids while on a buprenorphine/naloxone treatment [64], and another concluded that women and men respond differently to naloxone depending on the dose [65]. While we do not believe that male/female mice are akin to male/female humans (respectively), understanding the behavior and physiology of male and female mice provide a biological spectrum to understand the complexity across human populations.

### 4.5. Sleep deprivation one week after withdrawal result in changes in sleep on recovery day in males and females

Examining sleep behaviors following sleep deprivation allows us to analyze how our manipulations alter homeostatic sleep drive. Sleep deprivation, however, is also a stressor that many of us experience in our daily lives. Therefore, we wanted to observe how a 4-hour sleep deprivation early in the dark cycle would alter future sleep behavior. We performed these experiments 6 days after the final drug treatments because all mice returned to regular sleep cycles by day 3 of recovery (data not shown). We observed interesting qualitative differences between the sexes. On the day of sleep deprivation, we observed significant interactions and differences between groups that received naloxone (MN or SN) and those that did not (MS or SS) in male mice, but not female mice (**Fig. 7 & 8: A-C**). Interestingly, MS-male mice continued to sleep differently than MN and SN into the recovery day (**Fig. 7D-F)**. This reinforced the finding that spontaneous withdrawal and exacerbated withdrawal result in different experiences. All groups, in both sexes, slept more than they did at baseline **(Fig. 9 & 10: C,F)**. However, male and female mice who had received naloxone, regardless of morphine status, slept most similarly to their baselines on the deprivation day **(Fig. 9 & 10C)**. This result was surprising because we would have expected that the SS mice would sleep most similarly to baseline.

These data suggest that the stress of repeated saline injections profoundly affects sleep behavior, and should be considered when considering a saline injection as a control. Together, these changes might indicate that stressors occurring during abstinence impact male and female mice differently and that females may be more sensitive to the type of withdrawal experienced, whereas males seem more sensitive to the interaction between the morphine and naloxone. We know that stress is often a driving force behind relapse to drug-taking behavior and preclinical data implicates noradrenergic signaling in the extended amygdala in reinstatement [66–68]. Sleep, stress, and opioid signaling overlap in these extended amygdala circuits, and all utilize noradrenergic signaling. Additional studies are needed to determine the impact of these specific circuits on sleep behavior in withdrawal, but they present a space for interesting sex differences.

Our main finding is that male mice are more sensitive to morphine withdrawal-induced acute sleep disturbances than female mice, and female mice might be more “resilient” in response to future sleep-related stressors. These effects are notable as both male and female mice show behavioral adaptations well into protracted abstinence from this same paradigm [5], although those adaptations manifest in distinct ways. Continued use of both sexes of animals is critical to further understanding of how these pharmacological challenges manifest in alterations in physiology across neural systems and behavior.

### 4.6. Limitations

Sleep patterns are very specific to individuals due to the drive of social and environmental pressures in addition to genetic and biological drivers. As such, we see significant variability between subjects. This variability can often make it hard to identify treatment effects. There can also be a lot of variability due to environmental changes. We feel confident that the effects we report are accurate because our results were consistent across cohorts and cohorts consisted of appropriate controls run in parallel. Additionally, the within-subjects’ normalization of the data (Fig. 5, 6, 9, 10) confirms the overall trends we see in between-subjects’ comparisons. It is important to note that C57BL/6J mice, and many other inbred strains, are deficient in their melatonin production, a hormone important in regulating sleep [69–73]. Future studies would benefit from the inclusion of outbred strains or a panel of inbred strains.

### 4.7. Conclusions

Our study here is foundational for our future work examining sleep and opioids, and further validates the 3-day morphine withdrawal paradigm as one model for OUD that captures elements of the human condition in a mouse model where signaling and circuits can be assessed.

## Supporting information

supplemental statistics

## Funding

### The author(s) disclosed receipt of the following financial support for the research, authorship, and/or publication of this article

This work was supported by the Foundation of Hope; UNC-Pharmacological Sciences T32 [5T32GM135095]; National Institute of Drug Abuse [R01DA049261]; National Institute of Drug Abuse [NRSA/ F31DA05621].

## Declaration of Conflicting Interests

The authors M. Bedard and Dr. McElligott are sub-contracted by EpiCypher® on a SBIR grant unrelated to the work completed in this article.

## Author Contributions

MLB conceptualized experiments, conducted experiments, analyzed data, and wrote manuscript. JSL conducted experiments and analyzed data. IMB, AT, and PJP conducted experiments. LT and GD advised on experiments and provided feedback on the manuscript. ZAM conceptualized experiments, analyzed data, and wrote manuscript.

## Notes

### Competing Interest Statement

The authors have declared no competing interest.

### Summary of Updates

We have altered our analysis of the groups and largely combined groups to make more extensive cross group comparisons. While our overall interpretations have not changed dramatically, there have been extensive changes to the figures and analysis.

