## supplemental statistics for "Probing different paradigms of morphine withdrawal on sleep behavior in male and female C57BL/6J mice"

| Day | Time Frame | Source of Variation | % Time Asleep | | | Bout Length | | |
|---|---|---|---|---|---|---|---|---|
| | | | | F (DFn, DFd) | P value | | F (DFn, DFd) | P value |
| WD 1 | ZT 0-11 | ZT | **** | F (6.603, 317.0) = 32.88 | P<0.0001 | **** | F (2, 96) = 42.90 | P<0.0001 |
| | | Naloxone v Saline (Inj 2) | * | F (1, 48) = 5.833 | P=0.0196 | ns | F (1, 48) = 0.08051 | P=0.7778 |
| | | Morphine v Saline (Inj 1) | *** | F (1, 48) = 14.53 | P=0.0004 | ns | F (1, 48) = 3.606 | P=0.0636 |
| | | ZT x Naloxone v Saline (Inj 2) | ns | F (11, 528) = 1.511 | P=0.1235 | ns | F (2, 96) = 0.1441 | P=0.8660 |
| | | ZT x Morphine v Saline (Inj 1) | **** | F (11, 528) = 6.808 | P<0.0001 | ** | F (2, 96) = 6.089 | P=0.0032 |
| | | Naloxone v Saline (Inj 2) x Morphine v Saline (Inj 1) | * | F (1, 48) = 4.559 | P=0.0379 | ns | F (1, 48) = 0.03440 | P=0.8536 |
| | | ZT x Naloxone v Saline (Inj 2) x Morphine v Saline (Inj 1) | ns | F (11, 528) = 1.475 | P=0.1367 | ns | F (2, 96) = 0.04217 | P=0.9587 |
| | ZT 12-23 | ZT | **** | F (8.284, 397.6) = 11.53 | P<0.0001 | * | F (2, 96) = 4.433 | P=0.0144 |
| | | Naloxone v Saline (Inj 2) | ns | F (1, 48) = 0.3938 | P=0.5333 | ns | F (1, 48) = 0.01655 | P=0.8982 |
| | | Morphine v Saline (Inj 1) | **** | F (1, 48) = 18.83 | P<0.0001 | * | F (1, 48) = 5.521 | P=0.0229 |
| | | ZT x Naloxone v Saline (Inj 2) | ns | F (11, 528) = 1.519 | P=0.1204 | * | F (2, 96) = 3.831 | P=0.0251 |
| | | ZT x Morphine v Saline (Inj 1) | ns | F (11, 528) = 0.8567 | P=0.5831 | ns | F (2, 96) = 0.6064 | P=0.5474 |
| | | Naloxone v Saline (Inj 2) x Morphine v Saline (Inj 1) | ns | F (1, 48) = 2.985 | P=0.0905 | ns | F (1, 48) = 0.01656 | P=0.8981 |
| | | ZT x Naloxone v Saline (Inj 2) x Morphine v Saline (Inj 1) | ns | F (11, 528) = 1.000 | P=0.4447 | ns | F (2, 96) = 0.1512 | P=0.8599 |
| WD 2 | ZT 0-11 | ZT | **** | F (6.440, 309.1) = 47.76 | P<0.0001 | **** | F (2, 96) = 91.61 | P<0.0001 |
| | | Naloxone v Saline (Inj 2) | ns | F (1, 48) = 0.002384 | P=0.9613 | ns | F (1, 48) = 0.0008469 | P=0.9769 |
| | | Morphine v Saline (Inj 1) | **** | F (1, 48) = 43.02 | P<0.0001 | * | F (1, 48) = 4.258 | P=0.0445 |
| | | ZT x Naloxone v Saline (Inj 2) | ns | F (11, 528) = 1.338 | P=0.1993 | ns | F (2, 96) = 0.07506 | P=0.9277 |
| | | ZT x Morphine v Saline (Inj 1) | **** | F (11, 528) = 10.30 | P<0.0001 | *** | F (2, 96) = 9.242 | P=0.0002 |
| | | Naloxone v Saline (Inj 2) x Morphine v Saline (Inj 1) | ns | F (1, 48) = 1.466 | P=0.2319 | ns | F (1, 48) = 1.539 | P=0.2207 |
| | | ZT x Naloxone v Saline (Inj 2) x Morphine v Saline (Inj 1) | ns | F (11, 528) = 0.8167 | P=0.6234 | ns | F (2, 96) = 1.750 | P=0.1793 |
| | ZT 12-23 | ZT | **** | F (7.476, 358.9) = 12.23 | P<0.0001 | * | F (2, 96) = 3.123 | P=0.0485 |
| | | Naloxone v Saline (Inj 2) | ns | F (1, 48) = 0.05682 | P=0.8126 | ns | F (1, 48) = 0.8994 | P=0.3477 |
| | | Morphine v Saline (Inj 1) | **** | F (1, 48) = 39.78 | P<0.0001 | ** | F (1, 48) = 8.457 | P=0.0055 |
| | | ZT x Naloxone v Saline (Inj 2) | ** | F (11, 528) = 2.758 | P=0.0017 | ns | F (2, 96) = 0.7229 | P=0.4879 |
| | | ZT x Morphine v Saline (Inj 1) | ns | F (11, 528) = 0.8326 | P=0.6073 | ** | F (2, 96) = 5.932 | P=0.0037 |
| | | Naloxone v Saline (Inj 2) x Morphine v Saline (Inj 1) | ns | F (1, 48) = 0.002108 | P=0.9636 | ns | F (1, 48) = 0.2122 | P=0.6471 |
| | | ZT x Naloxone v Saline (Inj 2) x Morphine v Saline (Inj 1) | ns | F (11, 528) = 0.6435 | P=0.7916 | ns | F (2, 96) = 0.3928 | P=0.6762 |

**Supplemental Table 1.** Results from 3-way ANOVAs run for Figure 3 (Males). Continued on next page.

| Day | Time Frame | Source of Variation | % Time Asleep | | | Bout Length | | |
|---|---|---|---|---|---|---|---|---|
| | | | | F (DFn, DFd) | P value | | F (DFn, DFd) | P value |
| WD 3 | ZT 0-11 | ZT | **** | F (7.401, 355.2) = 47.37 | P<0.0001 | **** | F (2, 96) = 68.75 | P<0.0001 |
| | | Naloxone v Saline (Inj 2) | ns | F (1, 48) = 0.6542 | P=0.4226 | ns | F (1, 48) = 0.07274 | P=0.7885 |
| | | Morphine v Saline (Inj 1) | **** | F (1, 48) = 32.23 | P<0.0001 | ns | F (1, 48) = 0.2943 | P=0.5900 |
| | | ZT x Naloxone v Saline (Inj 2) | ns | F (11, 528) = 1.462 | P=0.1422 | ns | F (2, 96) = 1.116 | P=0.3319 |
| | | ZT x Morphine v Saline (Inj 1) | **** | F (11, 528) = 9.142 | P<0.0001 | ** | F (2, 96) = 6.753 | P=0.0018 |
| | | Naloxone v Saline (Inj 2) x Morphine v Saline (Inj 1) | ns | F (1, 48) = 0.6540 | P=0.4227 | ns | F (1, 48) = 0.09764 | P=0.7560 |
| | | ZT x Naloxone v Saline (Inj 2) x Morphine v Saline (Inj 1) | *** | F (11, 528) = 3.325 | P=0.0002 | ns | F (2, 96) = 1.415 | P=0.2478 |
| | ZT 12-23 | ZT | **** | F (7.988, 383.4) = 11.57 | P<0.0001 | * | F (2, 96) = 4.684 | P=0.0115 |
| | | Naloxone v Saline (Inj 2) | ns | F (1, 48) = 1.189 | P=0.2809 | ns | F (1, 48) = 1.120 | P=0.2952 |
| | | Morphine v Saline (Inj 1) | **** | F (1, 48) = 48.36 | P<0.0001 | *** | F (1, 48) = 13.54 | P=0.0006 |
| | | ZT x Naloxone v Saline (Inj 2) | **** | F (11, 528) = 3.590 | P<0.0001 | ns | F (2, 96) = 0.8332 | P=0.4378 |
| | | ZT x Morphine v Saline (Inj 1) | ns | F (11, 528) = 1.624 | P=0.0883 | ns | F (2, 96) = 1.104 | P=0.3357 |
| | | Naloxone v Saline (Inj 2) x Morphine v Saline (Inj 1) | ns | F (1, 48) = 1.493 | P=0.2278 | ns | F (1, 48) = 0.02057 | P=0.8866 |
| | | ZT x Naloxone v Saline (Inj 2) x Morphine v Saline (Inj 1) | ns | F (11, 528) = 0.8536 | P=0.5863 | ns | F (2, 96) = 0.1453 | P=0.8650 |
| Recovery | ZT 0-11 | ZT | **** | F (7.556, 362.7) = 9.831 | P<0.0001 | **** | F (2, 96) = 11.22 | P<0.0001 |
| | | Naloxone v Saline (Inj 2) | ns | F (1, 48) = 0.4112 | P=0.5244 | ns | F (1, 48) = 2.500 | P=0.1204 |
| | | Morphine v Saline (Inj 1) | ** | F (1, 48) = 7.455 | P=0.0088 | ns | F (1, 48) = 2.283 | P=0.1374 |
| | | ZT x Naloxone v Saline (Inj 2) | ns | F (11, 528) = 1.554 | P=0.1090 | * | F (2, 96) = 4.620 | P=0.0121 |
| | | ZT x Morphine v Saline (Inj 1) | ** | F (11, 528) = 2.847 | P=0.0012 | ns | F (2, 96) = 0.1033 | P=0.9020 |
| | | Naloxone v Saline (Inj 2) x Morphine v Saline (Inj 1) | ns | F (1, 48) = 1.310 | P=0.2581 | ns | F (1, 48) = 0.1004 | P=0.7528 |
| | | ZT x Naloxone v Saline (Inj 2) x Morphine v Saline (Inj 1) | ns | F (11, 528) = 0.5828 | P=0.8435 | ns | F (2, 96) = 0.3692 | P=0.6923 |
| | ZT 12-23 | ZT | **** | F (7.449, 357.5) = 17.41 | P<0.0001 | **** | F (2, 96) = 10.70 | P<0.0001 |
| | | Naloxone v Saline (Inj 2) | ns | F (1, 48) = 3.575 | P=0.0647 | ns | F (1, 48) = 0.5526 | P=0.4609 |
| | | Morphine v Saline (Inj 1) | *** | F (1, 48) = 14.32 | P=0.0004 | ns | F (1, 48) = 0.2325 | P=0.6319 |
| | | ZT x Naloxone v Saline (Inj 2) | **** | F (11, 528) = 3.583 | P<0.0001 | ns | F (2, 96) = 3.044 | P=0.0523 |
| | | ZT x Morphine v Saline (Inj 1) | * | F (11, 528) = 1.861 | P=0.0419 | ns | F (2, 96) = 0.9067 | P=0.4073 |
| | | Naloxone v Saline (Inj 2) x Morphine v Saline (Inj 1) | ns | F (1, 48) = 1.208 | P=0.2773 | ns | F (1, 48) = 0.3339 | P=0.5660 |
| | | ZT x Naloxone v Saline (Inj 2) x Morphine v Saline (Inj 1) | * | F (11, 528) = 2.132 | P=0.0169 | ns | F (2, 96) = 0.3023 | P=0.7398 |

**Supplemental Table 1 Cont.** Results from 3-way ANOVAs run for Figure 3 (Males).

| Day | Comparison | Source of Variation | Male | | | Female | | |
|---|---|---|---|---|---|---|---|---|
| | | | | F (DFn, DFd) | P value | | F (DFn, DFd) | P value |
| WD 1 | Light vs. Dark Cycle | Light Cycle X Treatment Group | **** | F (3, 48) = 10.51 | P<0.0001 | **** | F (3, 48) = 17.07 | P<0.0001 |
| | | Light Cycle | **** | F (1, 48) = 51.37 | P<0.0001 | **** | F (1, 48) = 68.42 | P<0.0001 |
| | | Treatment Group | ns | F (3, 48) = 0.6967 | P=0.5586 | ns | F (3, 48) = 1.030 | P=0.3876 |
| | | Mouse | ns | F (48, 48) = 0.1973 | P>0.9999 | ns | F (48, 48) = 1.028 | P=0.4624 |
| WD 2 | Light vs. Dark Cycle | Light Cycle X Treatment Group | **** | F (3, 48) = 10.51 | P<0.0001 | ** | F (3, 48) = 5.848 | P=0.0017 |
| | | Light Cycle | **** | F (1, 48) = 51.37 | P<0.0001 | **** | F (1, 48) = 41.09 | P<0.0001 |
| | | Treatment Group | ns | F (3, 48) = 0.6967 | P=0.5586 | ns | F (3, 48) = 0.01928 | P=0.9963 |
| | | Mouse | ns | F (48, 48) = 0.1973 | P>0.9999 | ns | F (48, 48) = 0.3715 | P=0.9996 |
| WD 3 | Light vs. Dark Cycle | Light Cycle X Treatment Group | *** | F (3, 48) = 7.344 | P=0.0004 | * | F (3, 48) = 3.065 | P=0.0367 |
| | | Light Cycle | **** | F (1, 48) = 375.0 | P<0.0001 | **** | F (1, 48) = 217.6 | P<0.0001 |
| | | Treatment Group | ns | F (3, 48) = 1.068 | P=0.3716 | ns | F (3, 48) = 0.2278 | P=0.8766 |
| | | Mouse | ns | F (48, 48) = 0.4093 | P=0.9988 | ns | F (48, 48) = 0.8745 | P=0.6780 |
| Recovery | Light vs. Dark Cycle | Light Cycle X Treatment Group | ** | F (3, 48) = 5.913 | P=0.0016 | ns | F (3, 48) = 1.302 | P=0.2846 |
| | | Light Cycle | **** | F (1, 48) = 388.1 | P<0.0001 | **** | F (1, 48) = 155.0 | P<0.0001 |
| | | Treatment Group | ns | F (3, 48) = 1.641 | P=0.1923 | ns | F (3, 48) = 0.9269 | P=0.4350 |
| | | Mouse | ns | F (48, 48) = 0.3647 | P=0.9997 | ns | F (48, 48) = 0.5523 | P=0.9789 |

**Supplemental Table 2.** Results from 2-way ANOVAs run for Figures 3 (Males) & 4 (Females).

| Day | Time Frame | Source of Variation | % Time Asleep | | | Bout Length | | |
|---|---|---|---|---|---|---|---|---|
| | | | | F (DFn, DFd) | P value | | F (DFn, DFd) | P value |
| WD 1 | ZT 0-11 | ZT | **** | F (6.527, 313.3) = 32.25 | P<0.0001 | **** | F (2, 96) = 68.74 | P<0.0001 |
| | | Naloxone v Saline (Inj 2) | ns | F (1, 48) = 2.960 | P=0.0918 | *** | F (1, 48) = 14.41 | P=0.0004 |
| | | Morphine v Saline (Inj 1) | ** | F (1, 48) = 12.01 | P=0.0011 | ns | F (1, 48) = 2.213 | P=0.1434 |
| | | ZT x Naloxone v Saline (Inj 2) | **** | F (11, 528) = 4.195 | P<0.0001 | ns | F (2, 96) = 1.584 | P=0.2104 |
| | | ZT x Morphine v Saline (Inj 1) | **** | F (11, 528) = 10.40 | P<0.0001 | *** | F (2, 96) = 8.255 | P=0.0005 |
| | | Naloxone v Saline (Inj 2) x Morphine v Saline (Inj 1) | ns | F (1, 48) = 1.057 | P=0.3090 | ns | F (1, 48) = 0.07612 | P=0.7838 |
| | | ZT x Naloxone v Saline (Inj 2) x Morphine v Saline (Inj 1) | * | F (11, 528) = 1.874 | P=0.0403 | ns | F (2, 96) = 0.4598 | P=0.6328 |
| | ZT 12-23 | ZT | **** | F (7.837, 376.2) = 6.822 | P<0.0001 | ns | F (2, 96) = 0.2529 | P=0.7771 |
| | | Naloxone v Saline (Inj 2) | ns | F (1, 48) = 1.034 | P=0.3144 | ns | F (1, 48) = 0.0004163 | P=0.9838 |
| | | Morphine v Saline (Inj 1) | * | F (1, 48) = 5.741 | P=0.0205 | ns | F (1, 48) = 0.3322 | P=0.5671 |
| | | ZT x Naloxone v Saline (Inj 2) | ** | F (11, 528) = 2.472 | P=0.0050 | * | F (2, 96) = 4.627 | P=0.0121 |
| | | ZT x Morphine v Saline (Inj 1) | ns | F (11, 528) = 0.6259 | P=0.8073 | ns | F (2, 96) = 0.1773 | P=0.8378 |
| | | Naloxone v Saline (Inj 2) x Morphine v Saline (Inj 1) | ns | F (1, 48) = 2.555 | P=0.1165 | ns | F (1, 48) = 0.5200 | P=0.4743 |
| | | ZT x Naloxone v Saline (Inj 2) x Morphine v Saline (Inj 1) | ns | F (11, 528) = 1.247 | P=0.2527 | ns | F (2, 96) = 1.117 | P=0.3314 |
| WD 2 | ZT 0-11 | ZT | **** | F (6.982, 335.1) = 43.40 | P<0.0001 | **** | F (2, 96) = 90.76 | P<0.0001 |
| | | Naloxone v Saline (Inj 2) | ns | F (1, 48) = 0.4944 | P=0.4854 | ** | F (1, 48) = 11.48 | P=0.0014 |
| | | Morphine v Saline (Inj 1) | **** | F (1, 48) = 23.48 | P<0.0001 | ns | F (1, 48) = 2.402 | P=0.1278 |
| | | ZT x Naloxone v Saline (Inj 2) | ns | F (11, 528) = 1.429 | P=0.1557 | * | F (2, 96) = 3.190 | P=0.0456 |
| | | ZT x Morphine v Saline (Inj 1) | **** | F (11, 528) = 8.847 | P<0.0001 | *** | F (2, 96) = 9.005 | P=0.0003 |
| | | Naloxone v Saline (Inj 2) x Morphine v Saline (Inj 1) | ns | F (1, 48) = 0.6630 | P=0.4195 | * | F (1, 48) = 5.379 | P=0.0247 |
| | | ZT x Naloxone v Saline (Inj 2) x Morphine v Saline (Inj 1) | ns | F (11, 528) = 1.670 | P=0.0768 | ns | F (2, 96) = 0.1996 | P=0.8194 |
| | ZT 12-23 | ZT | **** | F (7.838, 376.2) = 8.758 | P<0.0001 | ** | F (2, 96) = 6.523 | P=0.0022 |
| | | Naloxone v Saline (Inj 2) | ns | F (1, 48) = 2.536 | P=0.1178 | ns | F (1, 48) = 0.5165 | P=0.4758 |
| | | Morphine v Saline (Inj 1) | ** | F (1, 48) = 11.47 | P=0.0014 | ns | F (1, 48) = 3.488 | P=0.0679 |
| | | ZT x Naloxone v Saline (Inj 2) | ** | F (11, 528) = 2.713 | P=0.0021 | ns | F (2, 96) = 2.831 | P=0.0639 |
| | | ZT x Morphine v Saline (Inj 1) | * | F (11, 528) = 2.153 | P=0.0157 | ns | F (2, 96) = 2.162 | P=0.1207 |
| | | Naloxone v Saline (Inj 2) x Morphine v Saline (Inj 1) | ns | F (1, 48) = 1.775 | P=0.1891 | ns | F (1, 48) = 0.02225 | P=0.8821 |
| | | ZT x Naloxone v Saline (Inj 2) x Morphine v Saline (Inj 1) | ns | F (11, 528) = 0.6779 | P=0.7601 | ns | F (2, 96) = 0.8353 | P=0.4369 |

**Supplemental Table 3.** Results from 3-way ANOVAs run for Figure 4 (Females).

| Day | Time Frame | Source of Variation | % Time Asleep | | | Bout Length | | |
|---|---|---|---|---|---|---|---|---|
| | | | | F (DFn, DFd) | P value | | F (DFn, DFd) | P value |
| WD 3 | ZT 0-11 | ZT | **** | F (6.927, 332.5) = 38.51 | P<0.0001 | **** | F (2, 96) = 53.49 | P<0.0001 |
| | | Naloxone v Saline (Inj 2) | ns | F (1, 48) = 2.933 | P=0.0932 | ns | F (1, 48) = 0.1970 | P=0.6591 |
| | | Morphine v Saline (Inj 1) | **** | F (1, 48) = 35.23 | P<0.0001 | ns | F (1, 48) = 2.429 | P=0.1256 |
| | | ZT x Naloxone v Saline (Inj 2) | ns | F (11, 528) = 1.482 | P=0.1342 | ns | F (2, 96) = 0.4797 | P=0.6205 |
| | | ZT x Morphine v Saline (Inj 1) | **** | F (11, 528) = 8.247 | P<0.0001 | * | F (2, 96) = 3.128 | P=0.0483 |
| | | Naloxone v Saline (Inj 2) x Morphine v Saline (Inj 1) | ns | F (1, 48) = 0.3836 | P=0.5386 | ns | F (1, 48) = 0.09662 | P=0.7573 |
| | | ZT x Naloxone v Saline (Inj 2) x Morphine v Saline (Inj 1) | ns | F (11, 528) = 1.406 | P=0.1660 | ns | F (2, 96) = 1.122 | P=0.3299 |
| | ZT 12-23 | ZT | **** | F (7.938, 381.0) = 7.965 | P<0.0001 | ns | F (2, 96) = 2.581 | P=0.0809 |
| | | Naloxone v Saline (Inj 2) | ns | F (1, 48) = 1.031 | P=0.3149 | ns | F (1, 48) = 1.135 | P=0.2920 |
| | | Morphine v Saline (Inj 1) | *** | F (1, 48) = 17.15 | P=0.0001 | ** | F (1, 48) = 12.02 | P=0.0011 |
| | | ZT x Naloxone v Saline (Inj 2) | **** | F (11, 528) = 3.524 | P<0.0001 | * | F (2, 96) = 3.606 | P=0.0309 |
| | | ZT x Morphine v Saline (Inj 1) | ns | F (11, 528) = 1.191 | P=0.2900 | ns | F (2, 96) = 1.870 | P=0.1597 |
| | | Naloxone v Saline (Inj 2) x Morphine v Saline (Inj 1) | ns | F (1, 48) = 0.004277 | P=0.9481 | ns | F (1, 48) = 2.179 | P=0.1464 |
| | | ZT x Naloxone v Saline (Inj 2) x Morphine v Saline (Inj 1) | ns | F (11, 528) = 0.5996 | P=0.8297 | ns | F (2, 96) = 0.3130 | P=0.7320 |
| Recovery | ZT 0-11 | ZT | **** | F (7.953, 381.7) = 10.56 | P<0.0001 | **** | F (2, 96) = 17.64 | P<0.0001 |
| | | Naloxone v Saline (Inj 2) | ns | F (1, 48) = 0.1691 | P=0.6827 | ns | F (1, 48) = 0.5788 | P=0.4505 |
| | | Morphine v Saline (Inj 1) | * | F (1, 48) = 4.173 | P=0.0466 | ns | F (1, 48) = 1.001 | P=0.3221 |
| | | ZT x Naloxone v Saline (Inj 2) | ns | F (11, 528) = 1.764 | P=0.0573 | ns | F (2, 96) = 0.7465 | P=0.4767 |
| | | ZT x Morphine v Saline (Inj 1) | * | F (11, 528) = 2.095 | P=0.0192 | ns | F (2, 96) = 0.9070 | P=0.4072 |
| | | Naloxone v Saline (Inj 2) x Morphine v Saline (Inj 1) | ns | F (1, 48) = 0.3314 | P=0.5675 | ns | F (1, 48) = 1.222 | P=0.2744 |
| | | ZT x Naloxone v Saline (Inj 2) x Morphine v Saline (Inj 1) | ns | F (11, 528) = 0.6316 | P=0.8022 | ns | F (2, 96) = 0.3475 | P=0.7074 |
| | ZT 12-23 | ZT | **** | F (7.874, 377.9) = 6.556 | P<0.0001 | ** | F (2, 96) = 5.547 | P=0.0053 |
| | | Naloxone v Saline (Inj 2) | ns | F (1, 48) = 0.2984 | P=0.5874 | ns | F (1, 48) = 0.0004800 | P=0.9826 |
| | | Morphine v Saline (Inj 1) | * | F (1, 48) = 5.651 | P=0.0215 | ns | F (1, 48) = 2.926 | P=0.0936 |
| | | ZT x Naloxone v Saline (Inj 2) | *** | F (11, 528) = 3.096 | P=0.0005 | ns | F (2, 96) = 0.3765 | P=0.6872 |
| | | ZT x Morphine v Saline (Inj 1) | *** | F (11, 528) = 3.053 | P=0.0006 | ns | F (2, 96) = 2.789 | P=0.0665 |
| | | Naloxone v Saline (Inj 2) x Morphine v Saline (Inj 1) | ns | F (1, 48) = 0.5420 | P=0.4652 | ns | F (1, 48) = 1.702e-005 | P=0.9967 |
| | | ZT x Naloxone v Saline (Inj 2) x Morphine v Saline (Inj 1) | ns | F (11, 528) = 0.2297 | P=0.9955 | ns | F (2, 96) = 0.2606 | P=0.7711 |

**Supplemental Table 3 Cont.** Results from 3-way ANOVAs run for Figure 4 (Females).

| Day | Source of Variation | Difference from Baseline | F (DFn, DFd) | P value | Linear Regression | |
|---|---|---|---|---|---|---|
| WD 1 | ZT | **** | F (2.195, 105.3) = 119.2 | P<0.0001 | MN | Y = 5.670*X - 112.7 |
| | Naloxone v Saline (Inj 2) | ** | F (1, 48) = 8.671 | P=0.0050 | MS | Y = 7.160*X - 83.59 |
| | Morphine v Saline (Inj 1) | ** | F (1, 48) = 8.112 | P=0.0065 | SN | Y = 3.626*X - 44.43 |
| | ZT x Naloxone v Saline (Inj 2) | **** | F (23, 1104) = 5.751 | P<0.0001 | SS | Y = 7.336*X - 42.85 |
| | ZT x Morphine v Saline (Inj 1) | **** | F (23, 1104) = 6.526 | P<0.0001 | Are slopes different? | |
| | Naloxone v Saline (Inj 2) x Morphine v Saline (Inj 1) | ns | F (1, 48) = 0.004180 | P=0.9487 | F = 8.728. DFn = 3, DFd = 1240 | |
| | ZT x Naloxone v Saline (Inj 2) x Morphine v Saline (Inj 1) | ns | F (23, 1104) = 1.483 | P=0.0666 | P<0.0001 | |
| WD 2 | ZT | **** | F (2.571, 123.4) = 215.4 | P<0.0001 | MN | Y = 6.444*X - 112.7 |
| | Naloxone v Saline (Inj 2) | ns | F (1, 48) = 3.228 | P=0.0787 | MS | Y = 9.629*X - 128.5 |
| | Morphine v Saline (Inj 1) | **** | F (1, 48) = 22.34 | P<0.0001 | SN | Y = 5.755*X - 46.78 |
| | ZT x Naloxone v Saline (Inj 2) | **** | F (23, 1104) = 7.895 | P<0.0001 | SS | Y = 8.296*X - 51.18 |
| | ZT x Morphine v Saline (Inj 1) | **** | F (23, 1104) = 12.47 | P<0.0001 | Are slopes different? | |
| | Naloxone v Saline (Inj 2) x Morphine v Saline (Inj 1) | ns | F (1, 48) = 0.02543 | P=0.8740 | F = 13.87. DFn = 3, DFd = 1240 | |
| | ZT x Naloxone v Saline (Inj 2) x Morphine v Saline (Inj 1) | ns | F (23, 1104) = 0.3871 | P=0.9961 | P<0.0001 | |
| WD 3 | ZT | **** | F (2.543, 122.0) = 171.3 | P<0.0001 | MN | Y = 5.916*X - 111.8 |
| | Naloxone v Saline (Inj 2) | ** | F (1, 48) = 7.680 | P=0.0079 | MS | Y = 9.651*X - 123.8 |
| | Morphine v Saline (Inj 1) | **** | F (1, 48) = 21.19 | P<0.0001 | SN | Y = 5.039*X - 48.47 |
| | ZT x Naloxone v Saline (Inj 2) | **** | F (23, 1104) = 9.109 | P<0.0001 | SS | Y = 8.146*X - 47.73 |
| | ZT x Morphine v Saline (Inj 1) | **** | F (23, 1104) = 9.005 | P<0.0001 | Are slopes different? | |
| | Naloxone v Saline (Inj 2) x Morphine v Saline (Inj 1) | ns | F (1, 48) = 0.05109 | P=0.8221 | F = 20.14. DFn = 3, DFd = 1240 | |
| | ZT x Naloxone v Saline (Inj 2) x Morphine v Saline (Inj 1) | ns | F (23, 1104) = 0.3706 | P=0.9972 | P<0.0001 | |
| Recovery | ZT | **** | F (2.313, 111.0) = 66.09 | P<0.0001 | MN | Y = 2.102*X - 29.56 |
| | Naloxone v Saline (Inj 2) | * | F (1, 48) = 5.725 | P=0.0207 | MS | Y = 6.351*X - 43.89 |
| | Morphine v Saline (Inj 1) | ** | F (1, 48) = 11.20 | P=0.0016 | SN | Y = 2.908*X + 6.247 |
| | ZT x Naloxone v Saline (Inj 2) | **** | F (23, 1104) = 13.80 | P<0.0001 | SS | Y = 7.201*X - 24.77 |
| | ZT x Morphine v Saline (Inj 1) | **** | F (23, 1104) = 3.439 | P<0.0001 | Are slopes different? | |
| | Naloxone v Saline (Inj 2) x Morphine v Saline (Inj 1) | ns | F (1, 48) = 0.5358 | P=0.4677 | F = 42.36. DFn = 3, DFd = 1240 | |
| | ZT x Naloxone v Saline (Inj 2) x Morphine v Saline (Inj 1) | ns | F (23, 1104) = 0.2748 | P=0.9998 | P<0.0001 | |

**Supplemental Table 4.** Results from 3-way ANOVAs and linear regressions run for Figure 5 (Males).

| Day | Source of Variation | Difference from Baseline | | | Linear Regression | |
|---|---|---|---|---|---|---|
| | | | F (DFn, DFd) | P value | | |
| WD 1 | ZT | **** | F (2.187, 105.0) = 70.37 | P<0.0001 | MN | Y = 5.626*X - 114.2 |
| | Naloxone v Saline (Inj 2) | *** | F (1, 48) = 14.43 | P=0.0004 | MS | Y = 4.746*X - 70.69 |
| | Morphine v Saline (Inj 1) | ** | F (1, 48) = 8.350 | P=0.0058 | SN | Y = 3.792*X - 72.91 |
| | ZT x Naloxone v Saline (Inj 2) | **** | F (23, 1104) = 3.473 | P<0.0001 | SS | Y = 6.477*X - 27.04 |
| | ZT x Morphine v Saline (Inj 1) | **** | F (23, 1104) = 2.776 | P<0.0001 | Are slopes different? | |
| | Naloxone v Saline (Inj 2) x Morphine v Saline (Inj 1) | ns | F (1, 48) = 2.233 | P=0.1416 | F = 3.544. DFn = 3, DFd = 1240 | |
| | ZT x Naloxone v Saline (Inj 2) x Morphine v Saline (Inj 1) | * | F (23, 1104) = 1.623 | P=0.0321 | P=0.0142 | |
| WD 2 | ZT | **** | F (2.153, 103.4) = 94.20 | P<0.0001 | MN | Y = 6.107*X - 114.5 |
| | Naloxone v Saline (Inj 2) | * | F (1, 48) = 4.317 | P=0.0431 | MS | Y = 6.180*X - 99.84 |
| | Morphine v Saline (Inj 1) | *** | F (1, 48) = 16.20 | P=0.0002 | SN | Y = 5.682*X - 67.27 |
| | ZT x Naloxone v Saline (Inj 2) | *** | F (23, 1104) = 2.499 | P=0.0001 | SS | Y = 7.370*X - 44.95 |
| | ZT x Morphine v Saline (Inj 1) | **** | F (23, 1104) = 5.056 | P<0.0001 | Are slopes different? | |
| | Naloxone v Saline (Inj 2) x Morphine v Saline (Inj 1) | ns | F (1, 48) = 0.9032 | P=0.3467 | F = 1.217. DFn = 3, DFd = 1240 | |
| | ZT x Naloxone v Saline (Inj 2) x Morphine v Saline (Inj 1) | ns | F (23, 1104) = 0.5450 | P=0.9605 | P=0.3021 | |
| WD 3 | ZT | **** | F (1.842, 88.40) = 73.34 | P<0.0001 | MN | Y = 5.066*X - 108.8 |
| | Naloxone v Saline (Inj 2) | ** | F (1, 48) = 8.255 | P=0.0060 | MS | Y = 6.915*X - 97.36 |
| | Morphine v Saline (Inj 1) | *** | F (1, 48) = 17.75 | P=0.0001 | SN | Y = 5.665*X - 65.00 |
| | ZT x Naloxone v Saline (Inj 2) | **** | F (23, 1104) = 2.727 | P<0.0001 | SS | Y = 6.351*X - 28.39 |
| | ZT x Morphine v Saline (Inj 1) | **** | F (23, 1104) = 4.841 | P<0.0001 | Are slopes different? | |
| | Naloxone v Saline (Inj 2) x Morphine v Saline (Inj 1) | ns | F (1, 48) = 0.1939 | P=0.6616 | F = 2.829. DFn = 3, DFd = 1240 | |
| | ZT x Naloxone v Saline (Inj 2) x Morphine v Saline (Inj 1) | ns | F (23, 1104) = 0.4571 | P=0.9872 | P=0.0374 | |
| Recovery | ZT | **** | F (2.086, 100.1) = 28.04 | P<0.0001 | MN | Y = 1.393*X - 24.56 |
| | Naloxone v Saline (Inj 2) | ns | F (1, 48) = 3.375 | P=0.0724 | MS | Y = 4.575*X - 52.65 |
| | Morphine v Saline (Inj 1) | ** | F (1, 48) = 7.274 | P=0.0096 | SN | Y = 3.072*X - 24.53 |
| | ZT x Naloxone v Saline (Inj 2) | **** | F (23, 1104) = 4.728 | P<0.0001 | SS | Y = 5.313*X - 12.53 |
| | ZT x Morphine v Saline (Inj 1) | **** | F (23, 1104) = 2.963 | P<0.0001 | Are slopes different? | |
| | Naloxone v Saline (Inj 2) x Morphine v Saline (Inj 1) | ns | F (1, 48) = 1.348 | P=0.2513 | F = 15.99. DFn = 3, DFd = 1240 | |
| | ZT x Naloxone v Saline (Inj 2) x Morphine v Saline (Inj 1) | ns | F (23, 1104) = 1.128 | P=0.3058 | P<0.0001 | |

**Supplemental Table 5.** Results from 3-way ANOVAs and linear regressions run for Figure 6 (Females).

| Day | Time Frame | Source of Variation | % Time Asleep | | | Bout Length | | |
|---|---|---|---|---|---|---|---|---|
| | | | | F (DFn, DFd) | P value | | F (DFn, DFd) | P value |
| Sleep Deprivation (SD) | ZT 0-11 | ZT | **** | F (4.816, 231.2) = 166.3 | P<0.0001 | **** | F (1.760, 84.48) = 51.82 | P<0.0001 |
| | | Naloxone v Saline (Inj 2) | ns | F (1, 48) = 1.218 | P=0.2753 | ns | F (1, 48) = 3.360 | P=0.0730 |
| | | Morphine v Saline (Inj 1) | ns | F (1, 48) = 0.03224 | P=0.8583 | ns | F (1, 48) = 0.01441 | P=0.9049 |
| | | ZT x Naloxone v Saline (Inj 2) | *** | F (11, 528) = 3.161 | P=0.0004 | ** | F (2, 96) = 7.219 | P=0.0012 |
| | | ZT x Morphine v Saline (Inj 1) | *** | F (11, 528) = 3.176 | P=0.0003 | ns | F (2, 96) = 0.6536 | P=0.5224 |
| | | Naloxone v Saline (Inj 2) x Morphine v Saline (Inj 1) | ns | F (1, 48) = 0.1458 | P=0.7042 | ns | F (1, 48) = 0.1294 | P=0.7207 |
| | | ZT x Naloxone v Saline (Inj 2) x Morphine v Saline (Inj 1) | ns | F (11, 528) = 1.247 | P=0.2526 | ns | F (2, 96) = 0.7105 | P=0.4939 |
| | ZT 12-23 | ZT | **** | F (7.662, 367.8) = 15.99 | P<0.0001 | ** | F (1.824, 87.55) = 6.290 | P=0.0037 |
| | | Naloxone v Saline (Inj 2) | ns | F (1, 48) = 3.540 | P=0.0660 | * | F (1, 48) = 6.989 | P=0.0110 |
| | | Morphine v Saline (Inj 1) | ** | F (1, 48) = 8.332 | P=0.0058 | * | F (1, 48) = 4.591 | P=0.0372 |
| | | ZT x Naloxone v Saline (Inj 2) | ns | F (11, 528) = 1.579 | P=0.1010 | ns | F (2, 96) = 1.001 | P=0.3712 |
| | | ZT x Morphine v Saline (Inj 1) | ns | F (11, 528) = 0.9291 | P=0.5117 | ns | F (2, 96) = 1.472 | P=0.2346 |
| | | Naloxone v Saline (Inj 2) x Morphine v Saline (Inj 1) | ns | F (1, 48) = 1.276 | P=0.2643 | ns | F (1, 48) = 1.811 | P=0.1847 |
| | | ZT x Naloxone v Saline (Inj 2) x Morphine v Saline (Inj 1) | ns | F (11, 528) = 1.011 | P=0.4346 | ns | F (2, 96) = 1.280 | P=0.2828 |
| Recovery | ZT 0-11 | ZT | **** | F (6.393, 306.9) = 10.04 | P<0.0001 | **** | F (1.883, 90.38) = 13.53 | P<0.0001 |
| | | Naloxone v Saline (Inj 2) | * | F (1, 48) = 5.553 | P=0.0226 | * | F (1, 48) = 4.783 | P=0.0336 |
| | | Morphine v Saline (Inj 1) | ns | F (1, 48) = 1.606 | P=0.2112 | ns | F (1, 48) = 1.152 | P=0.2885 |
| | | ZT x Naloxone v Saline (Inj 2) | **** | F (11, 528) = 4.539 | P<0.0001 | ns | F (2, 96) = 0.1567 | P=0.8552 |
| | | ZT x Morphine v Saline (Inj 1) | ns | F (11, 528) = 0.8754 | P=0.5645 | ns | F (2, 96) = 0.8626 | P=0.4253 |
| | | Naloxone v Saline (Inj 2) x Morphine v Saline (Inj 1) | ns | F (1, 48) = 0.4359 | P=0.5123 | ns | F (1, 48) = 3.682 | P=0.0610 |
| | | ZT x Naloxone v Saline (Inj 2) x Morphine v Saline (Inj 1) | ns | F (11, 528) = 0.8189 | P=0.6212 | ns | F (2, 96) = 0.6875 | P=0.5053 |
| | ZT 12-23 | ZT | **** | F (8.120, 389.8) = 19.11 | P<0.0001 | ns | F (1.551, 74.45) = 1.917 | P=0.1630 |
| | | Naloxone v Saline (Inj 2) | ** | F (1, 48) = 7.211 | P=0.0099 | ns | F (1, 48) = 3.166 | P=0.0815 |
| | | Morphine v Saline (Inj 1) | ns | F (1, 48) = 0.6275 | P=0.4322 | ns | F (1, 48) = 0.2521 | P=0.6179 |
| | | ZT x Naloxone v Saline (Inj 2) | **** | F (11, 528) = 3.547 | P<0.0001 | ns | F (2, 96) = 0.5438 | P=0.5823 |
| | | ZT x Morphine v Saline (Inj 1) | ns | F (11, 528) = 1.510 | P=0.1237 | ns | F (2, 96) = 0.7596 | P=0.4706 |
| | | Naloxone v Saline (Inj 2) x Morphine v Saline (Inj 1) | ns | F (1, 48) = 0.4846 | P=0.4897 | ns | F (1, 48) = 0.5987 | P=0.4429 |
| | | ZT x Naloxone v Saline (Inj 2) x Morphine v Saline (Inj 1) | ns | F (11, 528) = 1.657 | P=0.0800 | ns | F (2, 96) = 0.09020 | P=0.9138 |

**Supplemental Table 6.** Results from 3-way ANOVAs run for Figure 7 (Males).

| Day | Time Frame | Source of Variation | % Time Asleep | | | Bout Length | | |
|---|---|---|---|---|---|---|---|---|
| | | | | F (DFn, DFd) | P value | | F (DFn, DFd) | P value |
| Sleep Deprivation (SD) | ZT 0-11 | ZT | **** | F (6.242, 299.6) = 117.3 | P<0.0001 | **** | F (1.794, 128.3) = 66.70 | P<0.0001 |
| | | Naloxone v Saline (Inj 2) | ns | F (1, 48) = 0.7912 | P=0.3782 | ns | F (1, 143) = 0.2101 | P=0.6474 |
| | | Morphine v Saline (Inj 1) | ns | F (1, 48) = 0.3384 | P=0.5635 | ns | F (1, 143) = 0.5140 | P=0.4746 |
| | | ZT x Naloxone v Saline (Inj 2) | * | F (11, 528) = 1.952 | P=0.0311 | * | F (2, 143) = 3.082 | P=0.0489 |
| | | ZT x Morphine v Saline (Inj 1) | *** | F (11, 528) = 2.945 | P=0.0008 | ns | F (2, 143) = 0.7253 | P=0.4859 |
| | | Naloxone v Saline (Inj 2) x Morphine v Saline (Inj 1) | ns | F (1, 48) = 0.05403 | P=0.8172 | ns | F (1, 143) = 1.259 | P=0.2638 |
| | | ZT x Naloxone v Saline (Inj 2) x Morphine v Saline (Inj 1) | ns | F (11, 528) = 0.5664 | P=0.8564 | ns | F (2, 143) = 0.6343 | P=0.5318 |
| | ZT 12-23 | ZT | **** | F (7.172, 344.3) = 8.873 | P<0.0001 | ns | F (1.729, 83.01) = 2.719 | P=0.0795 |
| | | Naloxone v Saline (Inj 2) | ns | F (1, 48) = 0.7982 | P=0.3761 | ns | F (1, 48) = 2.134 | P=0.1506 |
| | | Morphine v Saline (Inj 1) | ns | F (1, 48) = 1.991 | P=0.1646 | ns | F (1, 48) = 0.2515 | P=0.6183 |
| | | ZT x Naloxone v Saline (Inj 2) | * | F (11, 528) = 1.835 | P=0.0457 | ns | F (2, 96) = 1.299 | P=0.2777 |
| | | ZT x Morphine v Saline (Inj 1) | ns | F (11, 528) = 1.093 | P=0.3645 | ns | F (2, 96) = 0.4525 | P=0.6374 |
| | | Naloxone v Saline (Inj 2) x Morphine v Saline (Inj 1) | ns | F (1, 48) = 0.2512 | P=0.6185 | ns | F (1, 48) = 0.03437 | P=0.8537 |
| | | ZT x Naloxone v Saline (Inj 2) x Morphine v Saline (Inj 1) | ns | F (11, 528) = 0.7094 | P=0.7300 | ns | F (2, 96) = 1.576 | P=0.2121 |
| Recovery | ZT 0-11 | ZT | **** | F (7.580, 363.9) = 9.758 | P<0.0001 | **** | F (1.886, 90.54) = 28.06 | P<0.0001 |
| | | Naloxone v Saline (Inj 2) | ns | F (1, 48) = 0.6128 | P=0.4376 | ns | F (1, 48) = 0.4260 | P=0.5171 |
| | | Morphine v Saline (Inj 1) | ns | F (1, 48) = 0.9780 | P=0.3276 | ns | F (1, 48) = 0.1628 | P=0.6884 |
| | | ZT x Naloxone v Saline (Inj 2) | ns | F (11, 528) = 1.079 | P=0.3762 | ns | F (2, 96) = 0.6800 | P=0.5091 |
| | | ZT x Morphine v Saline (Inj 1) | ns | F (11, 528) = 1.336 | P=0.2004 | ns | F (2, 96) = 1.715 | P=0.1855 |
| | | Naloxone v Saline (Inj 2) x Morphine v Saline (Inj 1) | ns | F (1, 48) = 1.244 | P=0.2702 | ns | F (1, 48) = 0.6533 | P=0.4229 |
| | | ZT x Naloxone v Saline (Inj 2) x Morphine v Saline (Inj 1) | ns | F (11, 528) = 0.3211 | P=0.9813 | ns | F (2, 96) = 0.07131 | P=0.9312 |
| | ZT 12-23 | ZT | **** | F (7.469, 358.5) = 8.706 | P<0.0001 | ns | F (1.836, 90.88) = 0.9988 | P=0.3666 |
| | | Naloxone v Saline (Inj 2) | ns | F (1, 48) = 0.03837 | P=0.8455 | ns | F (1, 50) = 1.312 | P=0.2576 |
| | | Morphine v Saline (Inj 1) | ns | F (1, 48) = 0.03965 | P=0.8430 | ns | F (1, 50) = 0.8102 | P=0.3724 |
| | | ZT x Naloxone v Saline (Inj 2) | ** | F (11, 528) = 2.604 | P=0.0031 | ns | F (2, 99) = 0.5063 | P=0.6043 |
| | | ZT x Morphine v Saline (Inj 1) | ns | F (11, 528) = 0.5429 | P=0.8741 | ns | F (2, 99) = 0.06694 | P=0.9353 |
| | | Naloxone v Saline (Inj 2) x Morphine v Saline (Inj 1) | ns | F (1, 48) = 0.2821 | P=0.5978 | ns | F (1, 50) = 2.094 | P=0.1541 |
| | | ZT x Naloxone v Saline (Inj 2) x Morphine v Saline (Inj 1) | ns | F (11, 528) = 1.287 | P=0.2281 | ns | F (2, 99) = 0.7480 | P=0.476 |

**Supplemental Table 7.** Results from 3-way ANOVAs run for Figure 8 (Females).

|  | Day | Source of Variation |  | Difference from Baseline F (DFn, DFd) | P value | Linear Regression |  |
|---|---|---|---|---|---|---|---|
| Males | SD | ZT | **** | F (23, 1104) = 224.2 | P<0.0001 | MN | Y = 1.842*X - 0.5436 |
|  |  | Naloxone v Saline (Inj 2) | * | F (1, 48) = 6.362 | P=0.0150 | MS | Y = 5.519*X - 28.01 |
|  |  | Morphine v Saline (Inj 1) | ns | F (1, 48) = 0.3638 | P=0.5493 | SN | Y = 2.534*X + 19.35 |
|  |  | ZT x Naloxone v Saline (Inj 2) | **** | F (23, 1104) = 7.193 | P<0.0001 | SS | Y = 7.333*X - 21.22 |
|  |  | ZT x Morphine v Saline (Inj 1) | ns | F (23, 1104) = 1.060 | P=0.3854 | Are slopes different? |  |
|  |  | Naloxone v Saline (Inj 2) x Morphine v Saline (Inj 1) | ns | F (1, 48) = 0.06629 | P=0.7979 | F = 47.43. DFn = 3, DFd = 1240 |  |
|  |  | ZT x Naloxone v Saline (Inj 2) x Morphine v Saline (Inj 1) | ns | F (23, 1104) = 0.3242 | P=0.9990 | P<0.0001 |  |
|  | Recovery | ZT | **** | F (23, 1104) = 75.39 | P<0.0001 | MN | Y = 5.161*X - 112.7 |
|  |  | Naloxone v Saline (Inj 2) | ns | F (1, 48) = 1.769 | P=0.1898 | MS | Y = 7.472*X - 116.2 |
|  |  | Morphine v Saline (Inj 1) | * | F (1, 48) = 6.287 | P=0.0156 | SN | Y = 5.356*X - 111.4 |
|  |  | ZT x Naloxone v Saline (Inj 2) | **** | F (23, 1104) = 18.94 | P<0.0001 | SS | Y = 8.350*X - 117.5 |
|  |  | ZT x Morphine v Saline (Inj 1) | ** | F (23, 1104) = 1.978 | P=0.0040 | Are slopes different? |  |
|  |  | Naloxone v Saline (Inj 2) x Morphine v Saline (Inj 1) | ns | F (1, 48) = 8.580e-005 | P=0.9926 | F = 11.99. DFn = 3, DFd = 1240 |  |
|  |  | ZT x Naloxone v Saline (Inj 2) x Morphine v Saline (Inj 1) | ns | F (23, 1104) = 0.5992 | P=0.9318 | P<0.0001 |  |
| Females | SD | ZT | **** | F (23, 1104) = 83.06 | P<0.0001 | MN | Y = 4.122*X - 94.42 |
|  |  | Naloxone v Saline (Inj 2) | ns | F (1, 48) = 0.8451 | P=0.3625 | MS | Y = 4.668*X - 104.4 |
|  |  | Morphine v Saline (Inj 1) | * | F (1, 48) = 5.760 | P=0.0203 | SN | Y = 4.381*X - 85.79 |
|  |  | ZT x Naloxone v Saline (Inj 2) | * | F (23, 1104) = 1.619 | P=0.0327 | SS | Y = 6.920*X - 92.26 |
|  |  | ZT x Morphine v Saline (Inj 1) | * | F (23, 1104) = 1.612 | P=0.0340 | Are slopes different? |  |
|  |  | Naloxone v Saline (Inj 2) x Morphine v Saline (Inj 1) | ns | F (1, 48) = 1.634 | P=0.2073 | F = 6.071. DFn = 3, DFd = 1240 |  |
|  |  | ZT x Naloxone v Saline (Inj 2) x Morphine v Saline (Inj 1) | ns | F (23, 1104) = 0.6043 | P=0.9285 | P=0.0004 |  |
|  | Recovery | ZT | **** | F (23, 1104) = 24.84 | P<0.0001 | MN | Y = 1.916*X - 4.235 |
|  |  | Naloxone v Saline (Inj 2) | ns | F (1, 48) = 0.2392 | P=0.6270 | MS | Y = 2.453*X - 27.10 |
|  |  | Morphine v Saline (Inj 1) | * | F (1, 48) = 6.203 | P=0.0163 | SN | Y = 2.584*X - 0.1093 |
|  |  | ZT x Naloxone v Saline (Inj 2) | **** | F (23, 1104) = 3.617 | P<0.0001 | SS | Y = 5.828*X - 6.804 |
|  |  | ZT x Morphine v Saline (Inj 1) | **** | F (23, 1104) = 2.715 | P<0.0001 | Are slopes different? |  |
|  |  | Naloxone v Saline (Inj 2) x Morphine v Saline (Inj 1) | ns | F (1, 48) = 2.760 | P=0.1031 | F = 10.63. DFn = 3, DFd = 1240 |  |
|  |  | ZT x Naloxone v Saline (Inj 2) x Morphine v Saline (Inj 1) | ns | F (23, 1104) = 1.257 | P=0.1865 | P<0.0001 |  |

**Supplemental Table 8.** Results from 3-way ANOVAs and linear regressions run for Figure 9 (Males) and Figure 10 (Females).
